# Cryo-EM Structure of Phospholipase Cε Defines N-terminal Domains and their Roles in Activity

**DOI:** 10.1101/2024.09.11.612521

**Authors:** Kadidia Samassekou, Elisabeth E. Garland-Kuntz, Vaani Ohri, Isaac J. Fisher, Satchal K. Erramilli, Kaushik Muralidharan, Livia M. Bogdan, Abigail M. Gick, Anthony A. Kossiakoff, Angeline M. Lyon

**Author notes:** To whom correspondence should be addressed: Angeline M. Lyon, the James Tarpo Jr. and Margaret Tarpo Department of Chemistry and the Department of Biological Sciences, Purdue University, 560 Oval Drive, West Lafayette, Indiana 47907.

## Abstract

Phospholipase Cε (PLCε) increases intracellular Ca^2+^ and protein kinase C (PKC) activity in the cardiovascular system in response to stimulation of G protein coupled receptors (GPCRs) and receptor tyrosine kinases (RTKs). The ability of PLCε to respond to these diverse inputs is due, in part, to multiple, conformationally dynamic regulatory domains. However, this heterogeneity has also limited structural studies of the lipase to either individual domains or its catalytic core. Here, we report the 3.9 Å reconstruction of the largest fragment of PLCε to date in complex with an antigen binding fragment (Fab). The structure reveals that PLCε contains a pleckstrin homology (PH) domain and four tandem EF hands, including subfamily-specific insertions and intramolecular interactions with the catalytic core. The structure, together with a model of the holoenzyme, suggest that part of the N-terminus and PH domain form a continuous surface that could engage cytoplasmic leaflets of the plasma and perinuclear membranes, contributing to activity. Functional characterization of this surface confirm it is critical for maximum basal and G protein-stimulated activities. This study provides new insights into the autoinhibited, basal conformation of PLCε and the first mechanistic insights into how it engages cellular membranes for activity.

## INTRODUCTION

Phospholipase Cs (PLCs) are highly conserved enzymes important for cell proliferation and survival^1^. All hydrolyze phosphatidylinositol-4,5-bisphosphate (PIP_2_) at the inner leaflet of the plasma membrane, producing inositol-1,4,5-trisphosphate (IP_3_) and diacylglycerol (DAG)^2^. IP_3_ stimulates the release of Ca^2+^ from intracellular stores, and the latter, together with DAG in the membrane, activate protein kinase C (PKC)^2^. PLCε is the largest PLC and its activity has been best characterized in the cardiovascular system. Under basal conditions, the lipase is autoinhibited in the cytoplasm. Stimulation of G_s_-or G_12/13_-coupled receptors leads to activation of the Rap1A or RhoA GTPases, respectively, which bind distinct sites on the lipase, leading to its translocation and activation at cell membranes^1, 3–5^. Rap1A·GTP stimulates PLCε activity at the perinuclear membrane, where phosphatidylinositol-4-phosphate (PI4P) is the substrate. PI4P hydrolysis activates a PKC-and protein kinase D (PKD)-dependent pathway that maximizes cardiac contractility^6, 7^. PLCε also activates Rap1A via its N-terminal CDC25 guanine nucleotide exchange factor (GEF) domain. This sustained, feed-forward activation loop is hypothesized to underlie the increased expression of pro-hypertrophic genes that leads to heart failure^8, 9^. This is consistent with the increased PLCε expression observed in patients with heart failure^10^. In contrast, RhoA·GTP activation of PLCε initiates a cardioprotective pathway, in which PIP_2_ dependent hydrolysis at the plasma membrane prevents mitochondrial apoptosis and cardiomyocyte death^11, 12^.

Almost all PLC enzymes share a highly conserved core defined by four domains: an N-terminal pleckstrin homology (PH) domain, four tandem EF hand repeats (EF1-4), a catalytic TIM barrel, and a C2 domain^1^ (**Fig. 1A**). In most subfamilies, these domains comprise the catalytic core required for lipid hydrolysis. However, in PLCε, EF hands 3/4 (EF3/4), the TIM barrel, C2 domain, and the Ras association (RA) 1 domain, form the catalytic core^13^. In this subfamily, the catalytic TIM barrel is stabilized through extensive intramolecular interactions between the RA1 domain, the EF3/4 hands, and the C2 domain^13^. Access to the active site is regulated in part by the X–Y linker, which splits the TIM barrel and physically prevents substrate binding in nearly all PLCs^13^. Lipase activity is further regulated by the C2-RA1 linker in PLCε, as mutation of the linker or its binding site on the catalytic core increase basal activity^13^.

**Figure 1.**
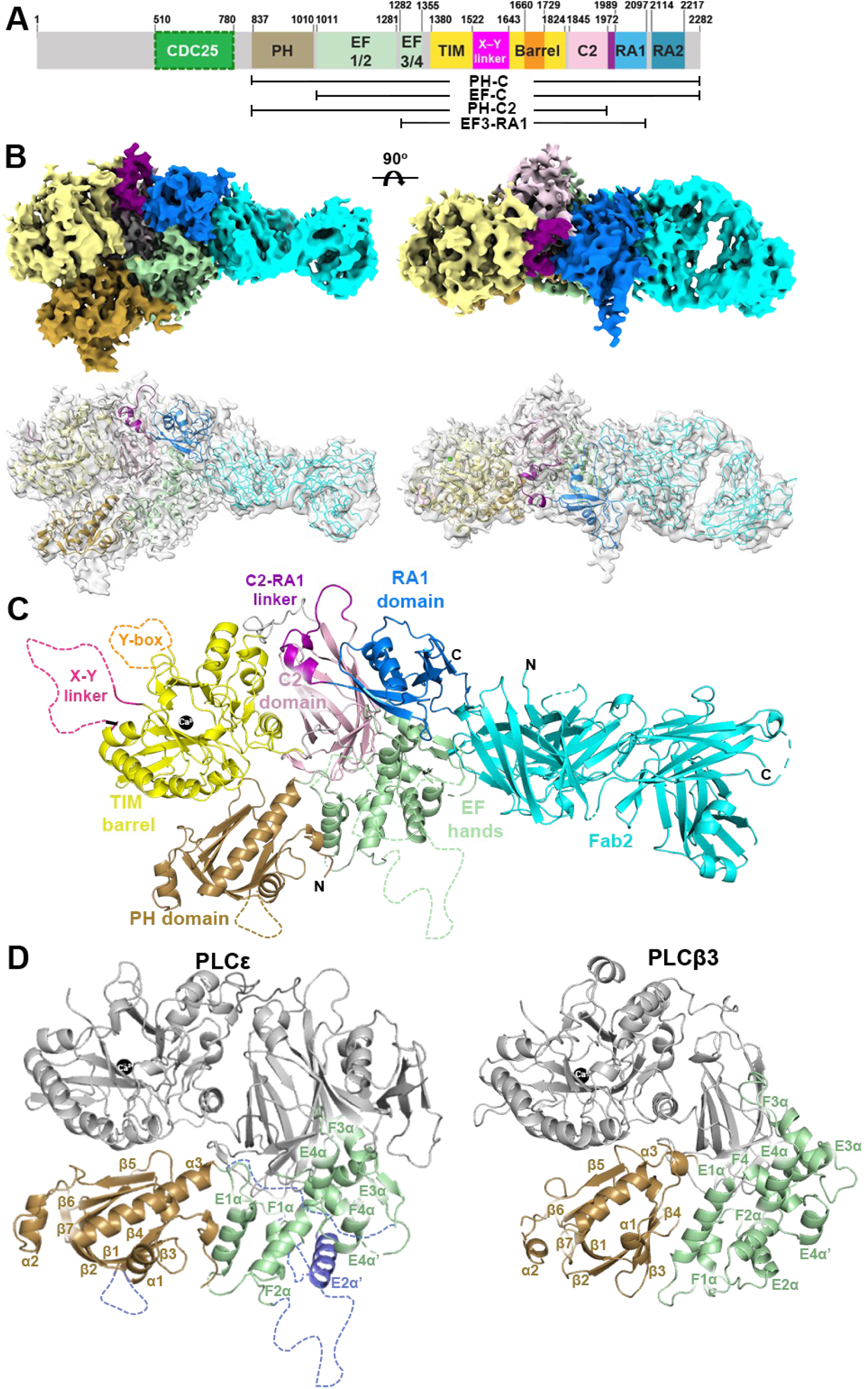
Cryo-EM reconstruction of the Fab2–PLCε PH-C complex. (**A**) Domain diagram of *R. norvegicus* PLCε. Numbers above correspond to experimentally determined domain boundaries, those predicted for the CDC25 domain are shown in dashed lines^17^. The Y-box (residues 1660-1729) is shown in orange and the C2-RA1 linker (residues 1972-1989) in purple. (**B**) (*Top*) Cryo-EM density map of the Fab2–PLCε PH-C complex at 3.9 Å resolution and (*bottom*) ribbon diagram fitted into the map. Domains in PLCε PH-C are colored as in Fig. 1A and Fab2 is shown in cyan. (**C**) Ribbon diagram of Fab2–PLCε PH-C. The active site Ca^2+^ is shown as a black sphere and disordered loops as dashed lines. The PLCε and Fab2 N-and C-termini are labeled N and C, respectively. (**D**) Ribbon diagrams of the PH-C2 domains of PLCε and PLCβ3 (PDB ID 3OHM)^25^. Secondary structural elements in the PH domains (wheat) and EF hands (pale green) are annotated based on PLCδ_29, 35_. Insertions in these regions unique to PLCε and the E2α’ helix are shown in blue.

The PLCε catalytic core is flanked by N-and C-terminal regulatory domains that contribute to membrane association, autoinhibition, and/or G protein-dependent activation. The first ∼800 residues of PLCε house the CDC25 domain, which is responsible for the GEF activity of the protein. This region is highly conserved across homologs (up to 90% identity), making it difficult to map domain boundaries. Within this region, residues 510-780 were predicted to form the minimum GEF domain, based on comparison to its closest homology, the Ras GEF Son-Of-Sevenless^8^. This region is separated from EF3/4, the start of the catalytic core, by ∼400 residues initially predicted to be a PH domain and EF hands 1/2. However, large insertions and minimal sequence identity to other PLCs (19% identity over this region)^14^ called these assignments into question. Biochemical studies of PLCε truncation variants that include this region demonstrated it promotes basal activity but has no impact on stability. This suggests that the region is flexibly connected to the EF3-RA1 catalytic core or their interactions are transient^15^. At the C-terminus, PLCε contains a second RA domain, RA2, that is flexibly connected to the catalytic core and serves as the primary binding site for Rap1A·GTP^9, 16^. Whether or how these regions interact with the PLCε catalytic core and/or the membrane to modulate activity is not known.

Here, we used cryo-electron microscopy (cryo-EM) single particle analysis (SPA) to determine the reconstruction of the largest fragment of PLCε to date in complex with an antigen binding fragment (Fab). The 3.9 Å reconstruction includes the ∼400 residues preceding the catalytic core and reveals this region does include PH and EF1/2 domains. The PH-EF1/2 module makes few contacts with the EF3-RA1 catalytic core, consistent with it being conformationally heterogeneous. The canonical membrane binding surface of the PLCε PH domain lacks the residues needed to bind substrate with high affinity and specificity, instead featuring conserved basic and hydrophobic residues. This surface is predicted to be extended by the CDC25 domain, which also makes extensive contacts with the PH domain, based on an AlphaFold2^17^ model of the holoenzyme. In cell-based assays, mutations disrupting the predicted interdomain interactions decreased activity, consistent with it being a functional interface. Similarly, mutations to the extended basic and hydrophobic surface formed by the CDC25 and PH domains decreased basal activity and Rap1A-and RhoA-dependent activation, suggesting the surface plays a general role in lipase activity. This study expands the known architecture of PLCε in its basal, autoinhibited conformation. Together with functional assays, it also provides the first insights into how this critical signaling enzyme engages membranes to carry out its roles in the cardiovascular system.

## RESULTS

### Generation of Fabs for PLCε structural studies

PLCε PH-C, an N-terminal truncation lacking the first 836 residues, was used for these studies because it can be purified in sufficient quantities for biophysical studies, retains sensitivity to G protein-dependent activation, and has been biochemically characterized^9, 13, 15^. It is also conformationally heterogenous, which limited prior attempts to obtain high-resolution structures^15^. PLCε PH-C, PLCε PH-C2 lacking the RA domains, and the PLCε EF3-RA1 catalytic core were used to generate a library of antigen binding fragments (Fabs) that decreased conformational heterogeneity and/or bound to a specific domain (**Fig. 1A**)^18–21^. After multiple rounds of selection and amplification starting from a naïve, fully synthetic phage display Fab library^22^, promising candidates were expressed, purified, and their binding affinity to each PLCε variant determined by surface plasmon resonance (SPR). Fab2 was selected for further study as it bound to PLCε PH-C and PLCε PH-C2 with K_D_ values of 4.4 nM and 22.9 nM, respectively, but did not bind PLCε EF3-RA1 (**Supporting Fig. 1A**).

### Cryo-EM SPA of the Fab2–PLCε PH-C Complex

A 1:1 Fab2–PLCε PH-C complex was isolated by size exclusion chromatography (SEC, **Supporting Fig. 1C**)^9, 15^. 7,016 micrographs were collected, and 4,090,458 particles picked using cryoSPARC^23^. Twelve 2D classes (690,149 particles) were selected based on visible secondary structure features and/or density characteristic of a Fab-bound protein. However, after refinement the initial map was highly anisotropic. This was corrected through iterative rounds of particle rebalancing and refinement to achieve a final structure with 3.9 Å resolution (**Supporting Fig. 2**). Local refinement was used to improve the resolution of PLCε PH-C or the Fab2– PLCε PH-C interface (**Supporting Fig. 3**), resulting in a final composite density map ranging from 3.0–7.0 Å resolution (**Fig. 1B**, **Supporting Fig. 4**). A homology model of Fab2 (PDB ID: 5BJZ)^24^ and the crystal structure of PLCε EF3-RA1 (PDB ID 6PMP)^13^ were rigid body fit into the final density map, revealing unmodeled density adjacent to the TIM barrel and C2 domains. Aligning the structure of the PLCβ3 catalytic core (PH-C2 domains, PDB ID 3OHM)^25^ with that of PLCε EF3-RA1 showed the PH and EF1/2 domains fit into this density. This region shares only 19% identity between PLCε and PLCβ3, and the former also contains large insertions. We therefore used AlphaFold2 to generate a preliminary initial model of PLCε for model building and refinement.

In the final reconstruction, the PLCε EF3-RA1 catalytic core is similar to the crystal structure (r.m.s.d. of 0.7 Å for 273 Cα residues) and the EF3-C2 domains in other PLCs. The largest differences are observed in the positions of EF3/4 and RA1 domains, and the C2-RA1 linker. This is likely due in part to the absence of lattice contacts in the cryo-EM reconstruction and binding of Fab2. The Fab binds to the EF3/4-RA1 interface, burying ∼2,200 Å^2^ surface area (**Fig. 1B,C**), resulting in EF3/4 moving ∼4 Å closer to the C2 domain while the C2-RA1 linker and RA1 domains move ∼5 Å and ∼7 Å away from the TIM barrel domain, respectively (**Fig. 1C**, **Supporting Figs. 5, 6**). No density is observed for the X–Y linker, Y-box, and RA2 domains, consistent with their conformational heterogeneity in solution^13^.

The PLCε PH and EF hands contain several subfamily-specific insertions, making them larger than their counterparts in other PLCs. The PH domain (residues 837-1010) maintains the canonical fold with the exception of a large, disordered loop (residues 892-929). Similar to the PLCβ PH domain, the membrane-binding surface of the PLCε PH domain lacks the residues required to bind PIP_2_ with high affinity and specificity^26–29^. Instead, this surface features solvent-exposed hydrophobic and basic residues, which in PLCβ engage the membrane via nonspecific interactions (**Supporting Figs. 7, 8**)^30–33^. Between the EF hands, EF1/2 (residues 1011-1281) is substantially larger than EF3/4 (residues 1282-1355) though both lack residues required for Ca^2+^ binding (**Supporting Figs. 5, 6**)^33, 34^. PLCε EF1/2 contains the E1α, F1α, and F2α helices, but lacks E2α, as do all PLCs except PLCβ^35^. This subdomain also contains a ∼140 amino acid insertion (residues 1050-1190) between the F1α and F2α helices of unknown function (**Fig. 1D**, **Supporting Fig. 7**). The PLCε EF3/4 module is highly homologous to other PLC enzymes, and includes the E3α, F3α, E4α, E4α’, and F4α helices. Interestingly, the E3α and F3αhelices are separated by a highly conserved ∼80 residue insertion (residues 1221-1300) that folds back on EF1/2, forms an a helix that packs against F1α and F2α, then re-enters the EF3/4 module. Because this helix interacts with EF1/2 the same way that the PLCβ E2α helix does, we refer to it as the E2α’ helix (**Fig. 1D**, **Supporting Fig. 7**).

### An intramolecular CDC25-PH domain interface contributes to activity

PLCε has higher basal activity than PLCε PH-C and other N-terminal truncation variants^13, 36^. We previously showed that the region we have now confirmed to be the PH domain increases basal activity and binding to PIP_2_-containing liposomes^15^. There must also be other domains within the first 836 residues that contribute to lipase activity^13, 36^. As this region has not yet been biochemically characterized, we turned to the AlphaFold2^17^ model of PLCε to identify structured regions that could be assessed (**Supporting Fig. 9**). No structure is predicted for the first ∼500 residues, despite the high degree of conservation (64-100% similarity). Only residues 510-780 are ordered, as they are predicted to correspond to the CDC25 domain, which has the same overall fold as the SOS GEF domain (**Supporting Fig. 9**)^37^. In the model, the G protein binding surface of the CDC25 domain is solvent-exposed, while the opposite face forms an extensive interface with the PH domain, burying ∼1,400 Å^2^ surface area (**Fig. 2A**, **Supporting Fig. 9**). The basic and hydrophobic membrane binding surface of the PH domain is extended by the CDC25 domain in this interaction, indicating the CDC25 domain may also contribute to lipase activity.

**Figure 2.**
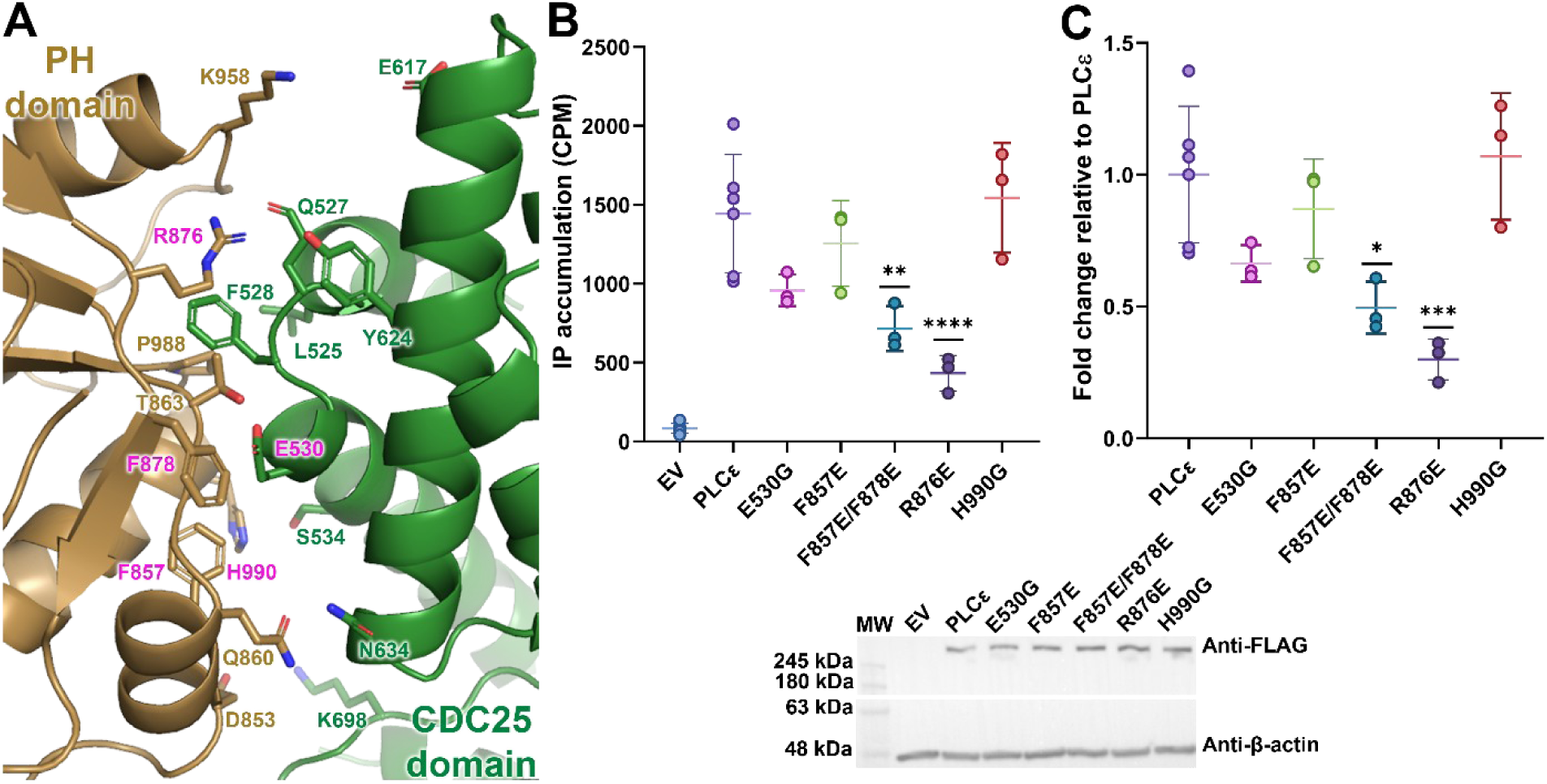
The CDC25 and PH domain form a functionally important interface. (**A**) The PLCε CDC25-PH domain interface predicted by AlphaFold2^17^. Residues in the interface are shown in sticks, with those labelled in magenta subject to mutagenesis. (**B**) Basal activity of the CDC25-PH domain point mutants was measured in cell-based assays. Each data point represents the average of one experiment in triplicate ± S.D., and EV refers to empty pCMV vector. (**C**) Fold change in basal activity relative to PLCε, where error bars correspond to S.D. All assays were performed from at least three independent transfections and analyzed using one-way ANOVA followed by Dunnett’s multiple comparisons test, ****p<0.0001, ***p=0.0006, **p=0.0022, *p=0.0100. A representative western blot of PLCε mutants expression is shown below (anti-FLAG at a 1:1000 dilution) with β-actin (anti-β-actin at a 1:1000 dilution) used as a loading control.

We first asked whether this predicted CDC25-PH domain interface was functionally relevant. Glutamate substitutions were introduced in PLCε and basal activity quantified in a cell-based inositol phosphate (IP_x_) accumulation assay^13^. PLCε F857 and F878 are conserved, solvent-exposed residues in the reconstruction of the Fab2–PLCε PH-C complex and thus poised to contribute to the CDC25-PH domain interface (**Fig. 2A**). PLCε F857E/F878E had ∼3-fold lower basal activity as compared to the wild-type protein (**Fig. 2B**).

PLCε R876 is poised to interact with Q527, F528, and Y624 in the CDC25 domain, and decreased activity ∼3.5 fold when mutated. Other mutations, including E530G in the CDC25 domain or F857E alone, also modestly decreased activity (**Fig. 2B, C**).

### Basic and hydrophobic residues on the CDC25-PH module are needed for maximum activity

PLCε must engage the membrane for its activity, but the membrane binding domain(s) have not been fully defined. The CDC25-PH module is a promising candidate, as removal of one or both domains decreases activity, and removal of the PH domain decreases binding to PIP_2_-containing liposomes^15^. Additionally, the CDC25-PH module in SOS was reported to contribute to membrane association of the enzyme^38^. To test this, site-directed mutagenesis was used to introduce glutamate residues to the surface of the CDC25 and/or PH domains in the background of PLCε, and their activity quantified^13^. PLCε EF-C, an N-terminally truncated variant (**Fig. 1A**), was used as a control as it has ∼6-fold lower basal activity^13, 15, 36^. Mutations to the CDC25 surface almost uniformly decreased activity. PLCε K618E/R619E had ∼3-fold lower activity than PLCε, while W607E, M615E, K652E, K655E, and M656E had ∼2-fold lower basal (**Fig. 3B, C**). On the PH domain, L873E, L928E, and H951E also caused modest decreases in basal activity (**Fig. 3B, C**).

**Figure 3.**
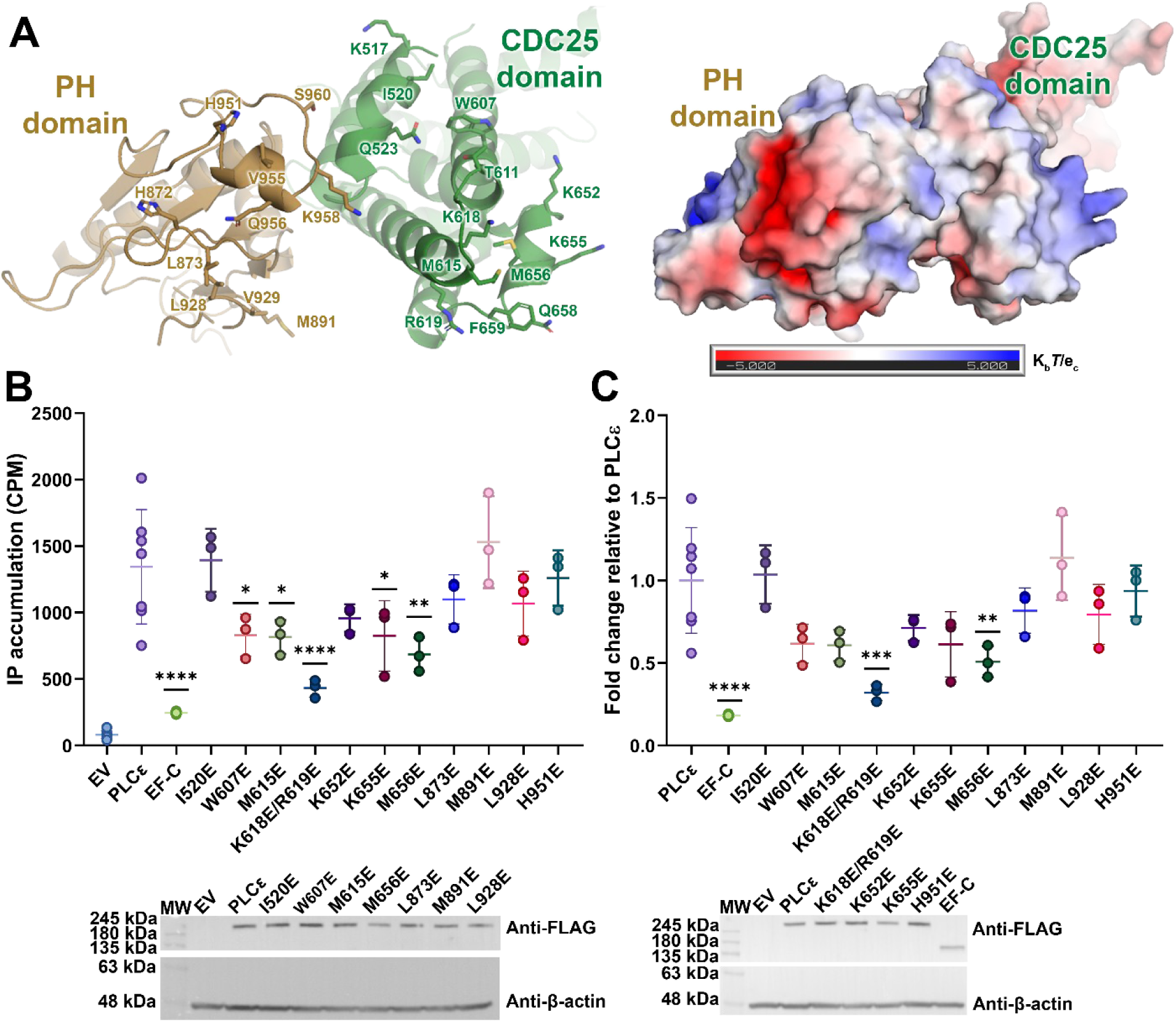
A basic and hydrophobic surface on the CDC25-PH domain module supports basal activity. (**A**) Ribbon diagram (*left*) and electrostatic surface (*right*) of the CDC25-PH domain module predicted by AlphaFold2^17^ and viewed through the membrane plane. Solvent-exposed residues on the surface are shown as sticks. (**B**) Basal activity of CDC25 and PH domain mutants was measured in cell-based assays. Each data point represents the average of one experiment performed in triplicate ± S.D., where EV is empty pCMV vector. (C) Fold change in activity ± S.D. relative to PLCε. Fold changes were calculated by dividing each mutant by PLCε. Assays were performed from at least three independent transfections in triplicate and analyzed using one-way ANOVA followed by Dunnett’s multiple comparisons test. ****p<0.0001, ***p=0.0002, **p≤0.0096, *p≤0.0365. Representative western blots showing PLCε mutant expression are shown below (anti-FLAG at a 1:1000 dilution) with β-actin (anti-β-actin at a 1:1000 dilution) used as a loading control.

If the CDC25-PH domain module serves as a general membrane interaction surface, we reasoned that maximum G protein-dependent activation should be similarly decreased. As the CDC25 and PH domains are dispensable for G protein-dependent activation, changes in activity are not due to defects in binding. We quantified RhoA-and Rap1A-dependent activation of PLCε, which occurs at the cytoplasmic leaflet of the plasma or perinuclear membrane, respectively^1, 3–5^. As the leaflets differ in their respective surface charge^1, 3–5^, it can also inform on the relative importance of the CDC25-PH domain residues at each subcellular site. For these studies, constitutively active Rap1A (Rap1A^Q63E^) and RhoA (RhoA^G14V^) were used^2, 39^. Co-transfection with Rap1A^Q63E^ increases PLCε activity ∼2.5-fold over basal^39^ (**Fig. 4A**). PLCε W607E, M615E, K618E/R619E, K655E, and M656E all had decreased maximum activity, up to ∼4-fold lower than PLCε. However, fold activation relative to each variant’s basal activity, was unchanged, as Rap1A^Q63E^ activated all PLCε variants ∼2-2.5 fold (**Fig. 4A**). RhoA^G14V^ increased PLCε activity ∼4.5-fold over basal (**Fig. 4B**). PLCε I520E, M656E, and M891E had similar maximum activities and fold activation relative to the wild-type PLCε. PLCε K618E/R619E had ∼2-fold lower maximum activity but maintained the same ∼4.5-fold activation relative to its basal as the other PLCε variants (**Fig. 4B**).

**Figure 4.**
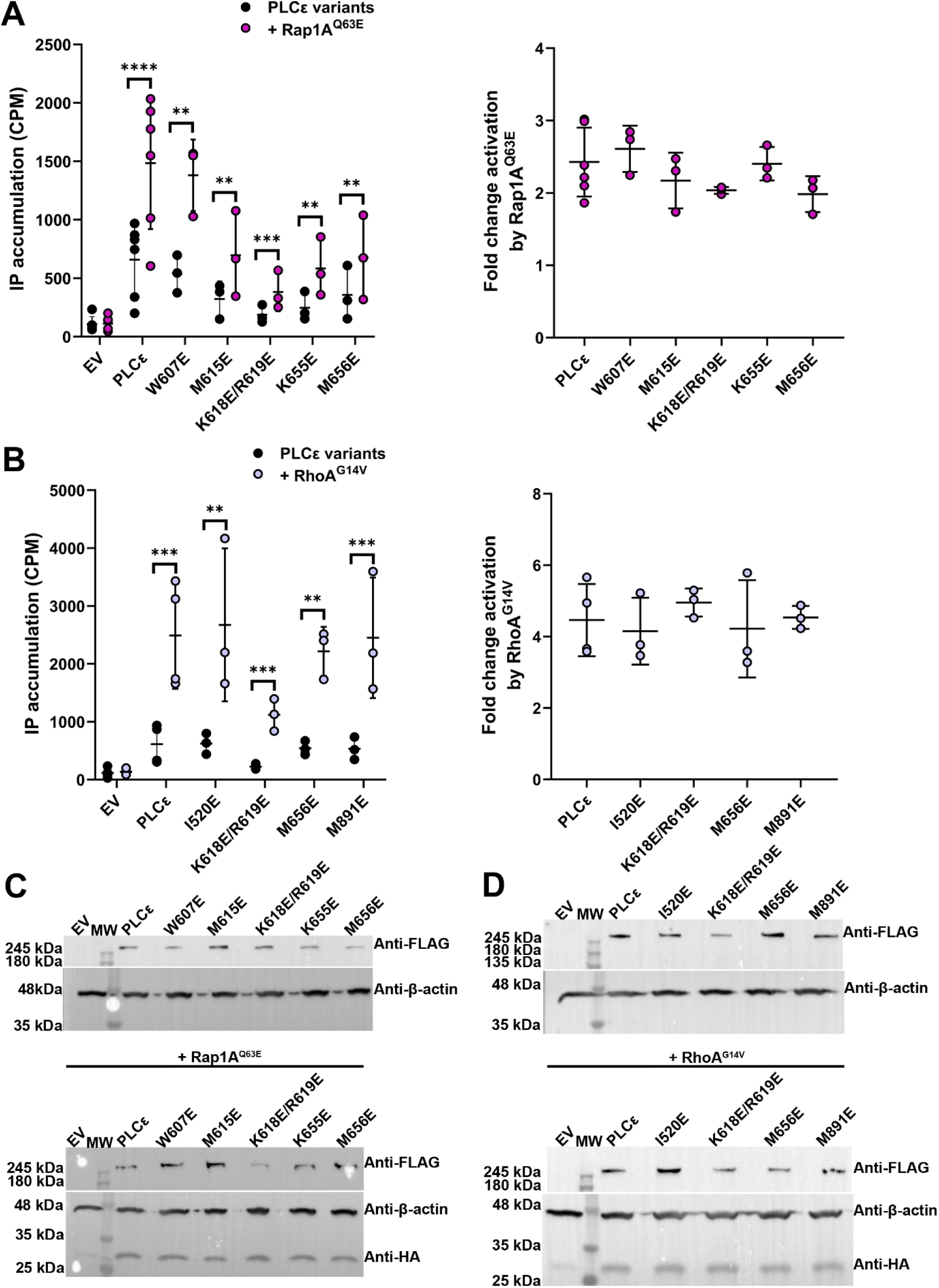
The CDC25-PH module surface contributes to Rap1A-and RhoA-dependent activation. Activation of the PLCε CDC25-PH domain mutants shown in Fig. 3 by constitutively active (**A**) Rap1A (Rap1A^Q63E^) and (**B**) RhoA (RhoA^G14V^) in cells. Each data point represents the average of one experiment performed in triplicate ± S.D. The fold change in activity is calculated by dividing the maximum G protein-stimulated activity of each PLCε variant by its corresponding basal activity. Assays were performed at least three times using independent transfections and analyzed using a ratio paired t-test, \*\*\*\**p*<0.0001, \*\**p*≤0.0005, \*\**p*≤0.0097. A one-way ANOVA and Dunnett’s multiple comparison test were used to test for differences in the fold-change activation of each mutant. Representative western blots showing PLCε variant (anti-FLAG at a 1:1000 dilution) and (**C**) Rap1A^Q63E^ or (D) RhoA^G14V^ (anti-HA at a 1:1000 dilution) expression are shown below, with EV corresponding to empty pCMV vector and β-actin (anti-β-actin at a 1:1000 dilution) used as a loading control.

## DISCUSSION

PLCε is required for normal cardiovascular function and its overexpression and/or sustained activation can lead to hypertrophy and heart failure^1, 4^. However, to validate PLCε as a potential therapeutic target requires a comprehensive understanding of its molecular architecture and regulation. Prior efforts to generate high-resolution structures of PLCε or fragments including its N-terminal regions failed due to conformational heterogeneity^15^. In the current study, we employed Fabs with the goal of stabilizing the variant and/or labelling specific domains. The use of Fab2 in the current study allowed us to determine the reconstruction of PLCε PH-C, the largest fragment to date (**Fig. 1**). The reconstruction allowed unambiguous identification and modeling of a PH domain and two tandem EF hands (EF1/2) preceding the catalytic core (**Fig. 1**). Fab2 does not bind the PH domain, instead it binds the EF hands and RA1 domain. Stabilization of these domain interfaces, especially those with EF1/2, likely restricts the movement of the PH domain, tethering it adjacent to the catalytic core.

The PLCε PH domain unites functional properties previously observed in other subfamilies. Like PLCδ, the PH domain is flexibly connected to the catalytic core, tethering it near the membrane^13, 15^. Like PLCβ, it lacks residues required to bind PIP_226-29_, instead interacting with the membrane through nonspecific interactions. Mutations to its surface are sufficient to decrease basal and G protein-stimulated activities of the holoenzyme in cells (**Figs. 3**, **4**). However, unlike other PLC enzymes, the PLCε PH domain is preceded by a CDC25 domain. These domains appear to form a single functional unit, as mutations within the interdomain surface decrease basal activity (**Fig. 2**, **Supporting Fig. 9**). This interaction also serves to expand the membrane interaction surface of the PH domain (**Fig. 2**). Mutations to residues on a solvent-exposed surface in the same plane as the active site decrease basal activity and maximum activation by Rap1A and RhoA, well-established regulators of the lipase (**Figs. 2-4**). Because the fold-activation by the G proteins was consistent across PLCε and all the point mutants, it is consistent with the basic and hydrophobic surface of the CDC25-PH module serving as a membrane association domain.

PLCε also diverges from other PLCs in its EF hands, especially EF1/2. While the EF hands are the least conserved region in the PH-C2 core observed in essentially all PLCs, PLCε EF1/2 shares only 25% identity in this region. EF1/2 features several large insertions, including a 140-residue loop within the EF1/2 module, that has not been functionally characterized. Interestingly, the fold of the EF1/2 is completed by the E2α’ helix, a subfamily-specific insertion that splits EF3/4 (**Fig. 1D**). Whether the EF hand insertions, or their intertwined structure, confer subfamily-specific regulation remains to be determined (**Fig. 1**, **Supporting Fig. 7**).

Based on this expanded understanding of PLCε architecture, we propose a model for its regulation in which formation and/or stabilization of intramolecular interactions primes the lipase to engage a membrane (**Fig. 5A**). For example, Fab2 binding the EF1/2-EF3/4-RA1 surface stabilizes the PH domain, and by extension the CDC25 domain, adjacent to the TIM barrel. Together with the active site in the TIM barrel, this allows the lipase to form multivalent interactions with a membrane surface. In the context of the holoenzyme, the N-terminal ∼500 residues may also contribute to membrane association and/or stabilization of intramolecular interactions, as their presence also contributes to activity^13, 15, 36^. G protein-dependent activation may utilize a similar mechanism. In the case of Rap1A, the PH and/or EF1/2 domains are required for activation and are stabilized in an extended conformation when the GTPase is bound^9^. The Rap1A-RA2 module (**Fig. 5B**), or a second molecule of Rap1A (**Fig. 5C**), could interact with EF1/2, stabilizing a more compact conformation of PLCε with increased activity^9^. Alternatively, the PLCε-specific insertions could directly contribute to G protein binding and/or stabilizing the activated state of the lipase at the membrane (**Fig. 5**). This study provides important structural insights into PLCε, including a more complete understanding of the autoinhibited, basal conformation of the lipase and insights into regions that contribute to membrane association. Further studies are needed to elucidate the structure and function of the PLCε N-terminus, and the roles of its numerous insertions. Identification of intra-and intermolecular interactions that occur upon binding the membrane and/or small GTPases will be needed to understand the roles of PLCε in healthy and diseased states.

**Figure 5.**
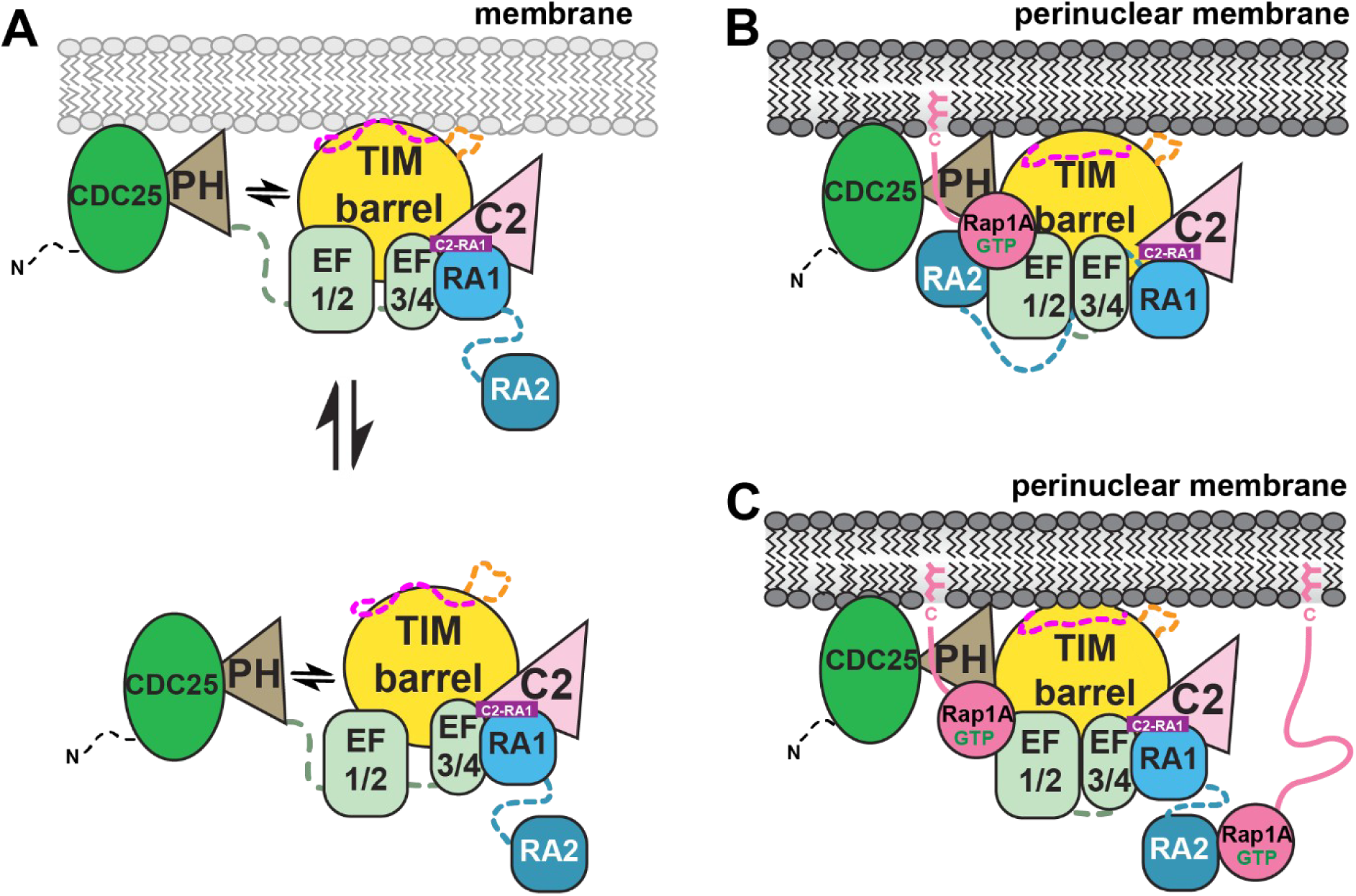
A model for PLCε regulation via stabilization of intramolecular interactions. **(A)** Under basal conditions, PLCε is autoinhibited in the cytoplasm by the X–Y and C2-RA1 linkers. The CDC25-PH module and EF1/2 are conformationally heterogenous in solution and interact transiently with the catalytic core. The lipase has basal activity and can interact with the membrane via the CDC25-PH module and TIM barrel. Dashed lines correspond to disordered loops. In response to stimulation of Gs-coupled receptors, Rap1A is activated and, at minimum, binds to the C-terminal RA2 domain. (**B**) The Rap1A-bound RA2 domain interacts with the PH domain and EF hands, stabilizing the intramolecular contacts between the CDC25-PH modules, EF hands, and TIM barrel. (**C**) Alternatively, a second Rap1A molecule binds to the CDC25-PH module and/or EF hands, stabilizing intramolecular interactions in the lipase. In either case, the multivalent interactions between PLCε, Rap1A, and the membrane help displace the X–Y linker, resulting in maximum activation at the perinuclear membrane.

## ACKNOWLEDGEMENTS

We thank Dr. John J. G. Tesmer and Dr. Charles Kuntz for technical and conceptual assistance. We thank Dr. Thomas Klose and Steve Wilson for assistance with cryo-EM data collection and computational work. This work is supported by the American Heart Association (23PRE1011279 to K.S.) and NHLBI (1R01HL141076-01 to A.M.L.). The content is solely the responsibility of the authors and does not necessarily represent the official views of the National Heart, Lung, and Blood Institute or the National Institutes of Health. Support from the Institute for Cancer Research Pilot Grant and Shared Resource Awards (P30CA023168 to A.M.L.), the Purdue Cryo-EM Facility, and the Chemical Genomics Facility at Purdue Institute for Drug Discovery are gratefully acknowledged. This work was supported in part by the Research Instrumentation Center in the Department of Chemistry.

## AUTHOR CONTRIBUTIONS

K.S., E.E.G.-K., and A.M.L. designed the experimental approach. S.K.E. and A.A.K. generated the Fabs and carried out binding assays. K.S. expressed and purified all proteins. Cryo-EM sample preparation and data collection were carried out by K.S., E.E.G.-K, and I.J.F. K.S processed the cryo-EM data, and K.S. and A.M.L. carried out refinement, and validation of the structure. K.S., K.M., L.M.B., and A.M.G. cloned PLCε variants for cell-based functional assays. K.S., V.O., and L.M.B. performed all activity assays. K.S. and A.M.L. wrote the manuscript.

## MATERIALS AND METHODS

### PLCε cloning, expression, and purification

A pCMV vector encoding *R. norvegicus* PLCε with a C-terminal FLAG tag (gift from A.V. Smrcka, U. Michigan) was used to generate all PLCε variants. Point mutants were introduced using the Q5 Site-Directed Mutagenesis Kit (New England BioLabs Inc) or In-Fusion cloning (Takara). PLCε PH-C (residues 837-2282), EF-C (residues 1038-2282), and EF3-RA1(residues 1284-2098) were subcloned into pFastBac HTA using QuickChange Site Directed Mutagenesis (Stratagene) or Q5 Site-Directed Mutagenesis Kit (New England BioLabs Inc) and expressed as N-terminally hexahistidine-tagged proteins. All plasmids were purified using the QIAprep Spin Miniprep Kit (Qiagen) and sequenced over the coding region.

PLCε PH-C, EF-C, and EF3-RA1 were expressed in baculovirus-infected Sf9 cells (RRID: CVCL_0549) at an MOI of ∼1 for 48 h at 27 °C with shaking at 120 rpm and harvested by centrifugation^5^. Pellets were flash frozen and stored at -80 °C until use. Frozen pellets were resuspended in lysis buffer containing 20 mM HEPES pH 8, 50 mM NaCl, 10 mM β-mercaptoethanol, 0.1 mM EGTA, 0.1 mM EDTA, two Roche EDTA-free protease inhibitor cocktail tablets (at one-third strength) and lysed by dounce on ice. The lysate was centrifuged for 1 h at 100,000 x *g* and the supernatant supplemented with NaCl and imidazole to final concentrations of 300 mM and 20 mM, respectively. The sample was loaded on a 5 mL HisTrap FF column (Cytiva) equilibrated with binding buffer (20 mM HEPES pH 8, 300 mM NaCl, 20 mM imidazole, 10 mM β-mercatopethanol, 0.1 mM EDTA, and 0.1 mM EGTA). After washing the column with 20 mL of binding buffer, the protein was eluted with a gradient of 0-100% binding buffer supplemented with 500 mM imidazole. Fractions containing protein were pooled, buffer exchanged, and concentrated with a 5-fold volume of MonoQ start buffer (20 mM HEPES pH 8, 50 mM NaCl, 2 mM DTT, 0.1 mM EDTA, and 0.1 mM EGTA) and injected onto a 1 mL MonoQ 5/50 GL column (Cytiva) pre-equilibrated with MonoQ start buffer. After washing the column with 10 mL of start buffer, protein was eluted with a gradient of 0-100% MonoQ start buffer supplemented to 500 mM NaCl. The protein was eluted, pooled, and concentrated to 1 mL prior to final purification over tandem Superdex 200 Increase columns (Cytiva) pre-equilibrated with S200 buffer (20 mM HEPES pH 8, 150 mM NaCl, 2 mM DTT, 0.1 mM EDTA, and 0.1 mM EGTA). Fractions corresponding to the purified protein were identified by SDS-PAGE, pooled, concentrated to ∼3-6 mg/mL, flash frozen in liquid nitrogen, and stored at -80 °C.

### Generation and validation of Fab2

Fab2 was generated by phage display using a protocol adapted from published methods^21, 40, 41^. PH-C, EF-C, and EF3-RA1 were chemically biotinylated overnight at 4°C using a 2-fold molar excess of NHS-PEG4-Biotin reagent (Thermo) in an amine-free buffer. Excess, unreacted label was removed by size-exclusion chromatography using a Superdex 200 column pre-equilibrated with running buffer (20 mM HEPES, pH 7.5, 150 mM NaCl). Fractions containing proteins were pooled, analyzed by SDS-PAGE, and biotinylation efficiency was quantified by test pull-downs using streptavidin paramagnetic particles (Promega). Four rounds of biopanning were performed by iteratively reducing the protein concentration from 200 to 10 nM in the final round. Initial validation was performed using single-point phage ELISA of individual clones from the final round phage pools. Positive clones were selected, subcloned into vector pRH2.2 (kind gift of Sachdev Sidhu), sequenced at the University of Chicago Comprehensive Cancer Center DNA Sequencing Facility, and expressed and purified. Epitope binning and kinetic analyses were performed using a MASS-1 instrument (Bruker) with a His-capture sensor chip (XanTec) as previously described^42^. A final concentration of 25-50 nM target protein, PLCε PH-C, EF-C, or EF3-RA1 were immobilized on the chip surface. Binding was quantified by surface plasmon resonance (SPR) using 1.5-200 nM Fab2 and running buffer (20 mM HEPES, pH 7.5, 150 mM NaCl, 50 mM EDTA, 0.05% Tween-20) in experimental and reference channels as a control. Sensorgrams were fitted to a one-to-one binding model and the K_D_ values were calculated.

### Fab2 expression and purification

Fab2 was expressed in *E. coli* BL21 cells via autoinduction in Terrific Broth supplemented with 0.4% glycerol, 0.01% glucose, 0.02% lactose, 1.25 mM MgSO_4_, and 100 µg/mL carbenicillin at 37 °C for 8 h, then overnight at 30 °C^21, 40, 41^. Cells were harvested by centrifugation and the pellets were flash frozen and stored at -80 °C until use. Pellets were resuspended in 35 mL lysis buffer (20 mM sodium phosphate buffer pH 7.4, 150 mM NaCl, and 1 mM PMSF), and lysed by sonication for 4 min on ice (1 sec on, 1 sec off pulses at 40% amplitude (QSONICA)). The lysate was incubated at 65 °C in a water bath for 30 min, cooled on ice for 15 minutes, and clarified by centrifugation at 8,200 x *g* for 30 minutes. The supernatant was filtered and applied to protein-L resin (GenScript) equilibrated with 5 column volume (CV) of running buffer (20 mM Tris, pH 7.5, 500 mM NaCl), and washed with 10 CV of running buffer. Fab2 was eluted with 5 CV of 0.1 M acetic acid. Fractions corresponding to the purified protein were identified by SDS-PAGE, pooled, buffer exchanged in 1 x phosphate-buffered saline (PBS) pH 7.4 and concentrated to ∼4-6 mg/mL. The protein was flash frozen in liquid nitrogen and stored at -80 °C.

### Fab2–PLCε PH-C complex formation and isolation

Fab2 and PLCε PH-C were buffer exchanged in complex buffer (20 mM HEPES, pH 8, 150 mM NaCl, 0.3 mM TCEP, 0.1 mM EGTA, 0.1 mM EDTA) prior to incubation at a 1.5:1 molar ratio of Fab2:PLCε PH-C on ice for 30 min in a final volume of 500 mL. in The sample was then applied to a single Superdex 200 column pre-equilibrated with complex buffer and fractions containing the complex confirmed by SDS-PAGE. These fractions were pooled and concentrated to 0.6 mg/mL and used to prepare cryo-EM samples.

### Cryo-EM sample preparation, data collection, and structure determination

0.6 mg/mL Fab2– PLCε PH-C complex was mixed with *n*-dodecyl-β-D-maltoside (DDM) (Anatrace) at a final concentration of 0.005%, and 3.5 mL of the sample applied to glow-discharged Quantifoil R1.2/1.3 300 mesh Cu grids, blotted at blot force 2 for 3 sec at 4.2 °C with 100% humidity and plunge-frozen in liquid ethane using a Vitrobot Mark IV (Thermo Fisher Scientific).

Cryo-EM data were collected on a Titan Krios G1 transmission electron microscope (Thermo Fisher Scientific) equipped with a K3 Summit direct electron detector (Gatan Inc) and Gatan Quantum GIF energy filter at a total dose of 53.4 e-/Å^2^ and a sampling of 0.539 Å/pixel on the specimen level. Patch motion correction and CTF estimation were performed using cryoSPARC^23^. An initial 5.0 Å cryo-EM reconstruction of Fab2–PLCε PH-C D1526-1546, a previously characterized variant lacking part of the X–Y linker, was used to generate 2D templates for particle picking in the Fab2–PLCε PH-C data set. 4,090,458 particles were extracted from 7,016 micrographs. Twelve 2D classes that showed well-ordered secondary structure and/or clear density for both proteins were selected and used to generate three *ab initio* reconstructions. The volumes with clear density for both Fab2 and PLCε PH-C were subjected to non-uniform refinement. However, the resulting model was anisotropic due to preferred orientation. This was corrected using 3D classification, 2D re-selection, and particle balancing, which resulted in Model *A* (46,170 particles). To increase the number of particles, more 2D classes were selected (1,471,782 particles) and used for *ab initio* reconstruction into four classes. One class represented Fab2–PLCε PH-C (Model *B*) and the other Fab2 alone (Model *C*). Models *A*, *B*, and *C* were then used in heterogeneous refinement to further separate the 1,471,782 particles. The best class with the most particles was further processed by 2D classification and particle balancing. Densities for PLCε PH-C and the Fab were improved using local refinement with different sized masks. The final composite map was generated by combining the two local refinement maps in PHENIX^43^. Structure modeling was performed by rigid body fitting the AlphaFold2^17^ model of PH-RA1 and a Fab2 model (PDB ID: 5BJZ)^24^ generated by I-Tasser^44^ into the density, followed by manual building in COOT^45^, and refinement and validation in PHENIX^43^.

### Cell-based [^3^H]-IPx accumulation assay

COS-7 cells (RRID: CVCL_0224) were plated at 100,000 cells per well in a 12-well plate in high-glucose Dulbecco’s Modified Eagle’s Medium (Corning) supplemented with 10% fetal bovine serum (FBS, Atlanta Biological), 1% Glutamax (Gibco), 1% penicillin-streptomycin (Corning), and incubated at 37 °C and 5% CO_213_. Cells were transfected the next day with 750 ng DNA encoding PLCε variants or empty vector, and 375 ng of Rap1A^Q63E^ or RhoA^G14V^, using FuGENE-6 (Promega). 24 h post-transfection, cells were washed once with serum-and inositol-free Ham’s F-10 media (Invitrogen), then incubated with Ham’s F-10 media supplemented with 1.5 mCi/well myo[2-^3^H(N)] inositol (Perkin Elmer/Revvity) for 16-18 h. Lithium chloride was added to the cells at a final concentration of 10 mM and incubated for 1 hour to inhibit the activity of inositol phosphatases. The media was aspirated, the cells washed once with ice-cold 1X PBS, and then lysed on ice for 30 min with 50 mM ice-cold formic acid. The lysates were applied to Dowex AG 1-X8 anion exchange columns (BioRad), washed twice with 50 mM formic acid, and once with 100 mM formic acid. [^3^H]-inositol phosphates were eluted with 1.2 M ammonium formate and 0.1 M formic acid into scintillation vials. Uniscint BD scintillation cocktail (National Diagnostics) was added in a 3:1 ratio, and samples counted on a scintillation counter. All experiments were performed at least three times in triplicate in independent transfections. Data was analyzed using a ratio paired t-test or one-way ANOVA followed by Dunnett’s multiple comparison test using GraphPad Prism version 10.2.3 for Windows GraphPad Software, Boston, Massachusetts USA, www.graphpad.com.

### Immunoblotting

Cells were lysed in sample buffer (100 mM Tris-HCl pH 6.8, 6% (w/v) sucrose, 2% (w/v) SDS, 5% (v/v) β-mercaptoethanol, and 0.02 % bromophenol blue) and incubated at 90 °C for 10 min before loading onto an SDS-PAGE gel^13^. The proteins were transferred to a 0.45µm polyvinylidene fluoride (PVDF) membrane (ThermoScientific) overnight. The next day, the membrane was blocked with 5% bovine serum albumin (Fisher) dissolved in 1X Tris-buffered saline and supplemented with 0.1% Tween-20 (TBST) for 1 hour, followed by incubation with the primary antibodies (1:1000), an anti-FLAG rabbit antibody (Cell Signaling Technology Cat# 14793, RRID:AB_2572291), anti-HA mouse antibody (Cell Signaling Technology Cat# 2367, RRID:AB_10691311), and anti-β-actin mouse antibody (Cell Signaling Technology Cat# 3700, RRID:AB_2242334) overnight. The membrane was washed three times with 1X TBST, then incubated with the anti-rabbit (Cell Signaling Technology Cat# 7074, RRID:AB_2099233) and anti-mouse secondary antibodies conjugated with HRP (Sigma-Aldrich Cat# 12-349, RRID:AB_390192) for 1 hour. The membrane was washed three times with 1X PBS, followed by addition of the ECL substrate (ThermoFisher) and imaged on a GeneGnome imager.

### Liposome-based activity assays

Inositol phosphate (IP) accumulation was measured using a modified version of the Cisbio IP-One assay^46, 47^. Briefly, 100 μM hen egg white phosphatidylethanolamine (PE) and 250 μM soy phosphatidylinositol phosphate (PI) (Avanti Polar Lipids) were mixed in chloroform, dried under nitrogen, and stored at -20 °C. The dried lipids were then resuspended in sonication buffer (50 mM HEPES, pH 7.4, 80 mM KCl, 3 mM EGTA, 1 mM DTT) and sonicated for 2 min. Reactions contained final concentrations of 2 ng PLCε PH-C, 1 mM Fab2, 50 mM HEPES, pH 7.4, 80 mM KCl, 16.7 mM NaCl, 0.83 mM MgCl_2_, 0.17 mM EDTA, 3 mM EGTA, 1 mM DTT, 1 mg/mL BSA, and 3 mM free Ca^2+^. Reactions were initiated by the addition of liposomes and transferred to a 30 °C heat block for 15 min. Control reactions lacked free Ca^2+^. Reactions were quenched with 5 mL quench buffer (50 mM HEPES, pH 7.4, 80 mM KCl, 3 mM EGTA, 1 mM DTT, 210 mM EGTA). 14 mL of each reaction was added to a 384-well white plate (Corning™ 3826), followed by 3 mL of reconstituted D2-labeled IP1 and anti-IP1 cryptate to measure the fluorescence. Plates were centrifuged for 2 minutes at 1000 x *g* at room temperature, incubated for 1 h, and read on a Synergy Neo2 HTS (Agilent) 620 and 665 nm. All experiments were performed at least three times in triplicate with protein from more than one purification. Total IP_1_ was quantified using a standard curve. Data was analyzed using a nonlinear regression (curve-fit) and converted to total IP_1_ (nM) followed by a two-tailed t-test using GraphPad Prism version 10.2.3 for Windows GraphPad Software, Boston, Massachusetts USA, www.graphpad.com.

## DATA AVAILABILITY

Data have been deposited to the Electron Microscopy Data Bank under accession codes EMD-44064, EMD-44065, EMD-44066, the Protein Data Bank under accession code 9B13, and the Electron Microscopy Public Archive under accession code EMPIAR-12020.

## SUPPORTING INFORMATION

**Table S1.** Cryo-EM data collection, refinement, and validation statistics.

|  |  |
| --- | --- |
| **Structure: Fab2–PLCε PH-C complex** |  |
| EMDB: EMD-44064, EMD-44065, EMD-44066, EMD-44067 |  |
| PDB: 9B13 |  |
| EMPIAR: EMPIAR-12020 |  |
| **Data collection** |  |
| Vitrification method | FEI Vitrobot |
| Microscope | Titan Krios |
| Magnification | 81000X |
| Voltage (kV) | 300 |
| Detector | GATAN K3 Summit |
| Total electron exposure (e<sup>-</sup>/Å<sup>2</sup>) | 53.75 |
| Number of frames | 40 |
| Defocus range (μm) | 0.7 – 2 |
| Pixel size (Å) | 0.539 |
| **Data processing** |  |
| Number of micrographs | 7,016 |
| Initial particle images (no.) | 1,471,782 |
| Final particle images (no.) | 147,120 |
| Symmetry imposed | C1 |
| Map resolution (Å) | 3.9 |
| FSC threshold | 0.143 |
| Map resolution range (Å) | 3.0–7.0 |
| **Refinement** |  |
| Model resolution (Å) | 4.1 |
| FSC threshold | 0.5 |
| Map sharpening *B* factor (Å<sup>2</sup>) | N/A |
| Map CC | 0.73 |
| Model composition |  |
| Non-hydrogen atoms | 10,014 |
| Protein residues | 1274 |
| Ligands | Ca<sup>2+</sup> |
| *B* factors (Å<sup>2</sup>, min/max/mean) |  |
| Protein | 55.93/202.18/111.98 |
| Ligand | 88.94/88.94/88.94 |
| R.m.s. deviations |  |
| Bond lengths (Å) | 0.004 |
| Bond angles (°) | 0.570 |
| Validation |  |
| MolProbity score | 2.46 |
| Clashscore | 10.01 |
| Poor rotamers | 3.72 |
| Ramachandran plot |  |
| Favored (%) | 91.49 |
| Allowed (%) | 8.43 |
| Disallowed (%) | 0.08 |

**Supporting Figure 1.**
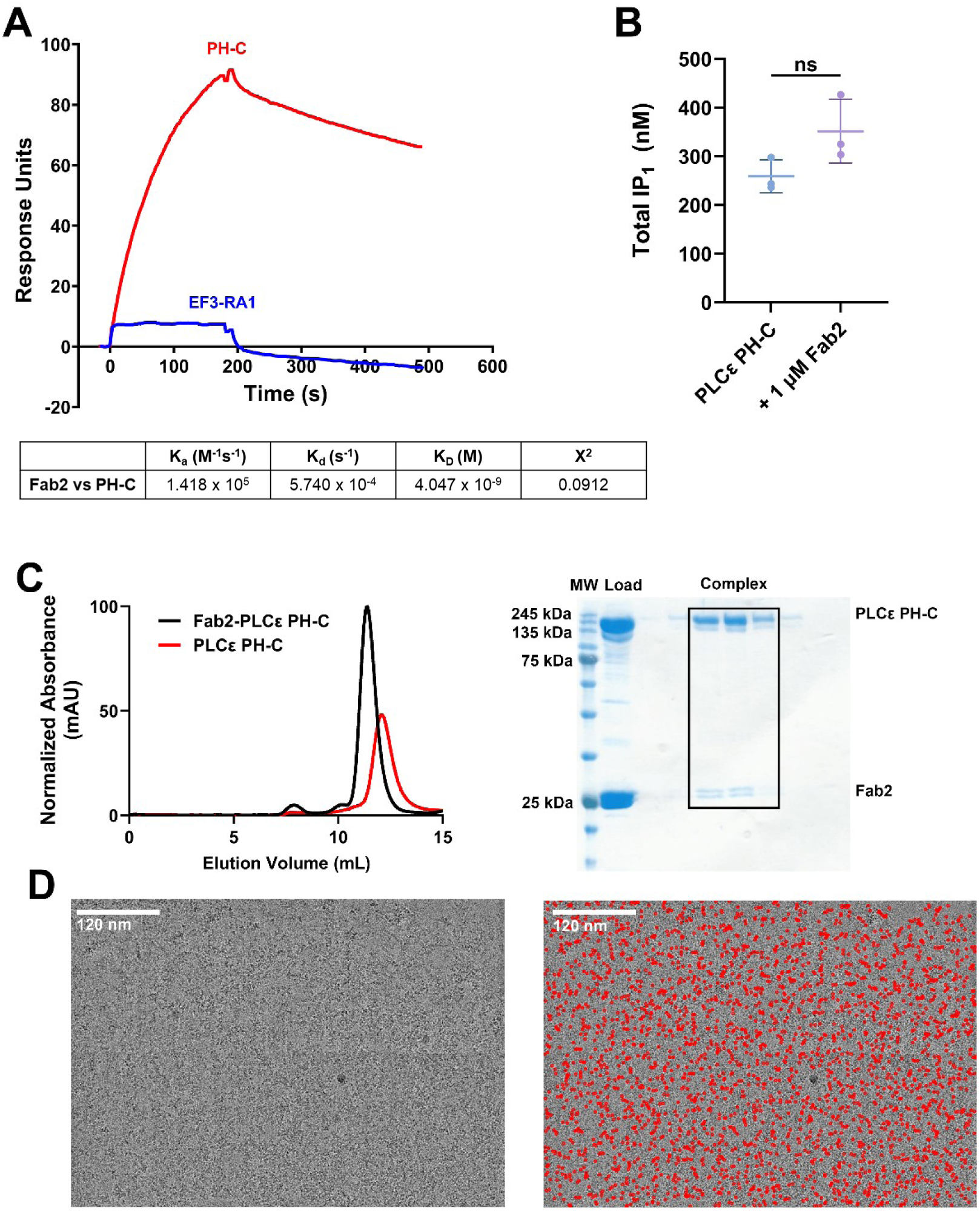
Fab2 characterization, complex isolation, and representative micrograph of the Fab2–PLCε PH-C complex. (**A**) Fab2 binds to a surface containing immobilized PLCε PH-C (red) but not EF3-RA1 (blue). Kinetic parameters are shown below. (**B**) Fab2 does not disrupt basal activity of PLCε PH-C in a liposome-based activity assay. PLCε PH-C was incubated with liposomes of 30% phosphatidylethanolamine (PE) and 70% phosphatidylinositol (PI), with or without 1 mM of Fab2. Total IP_1_ (nM) was calculated from a standard curve and analyzed using a two-tailed t-test (ns, not significant). Data represents the averages of three independent experiments carried out in triplicate using protein from at least two purifications. (**C**) (*Left*) Representative chromatogram of the Fab2–PLCε PH-C complex (black) and PH-C alone (red) on a Superdex 200 10/300 column. (*Right*) Coomassie-stained SDS-PAGE of the purified Fab2–PLCε PH-C complex (black box). (**D**) Representative micrograph of the Fab2–PLCε PH-C complex (*left*) with particles selected for analysis in red (*right*).

**Supporting Figure 2.**
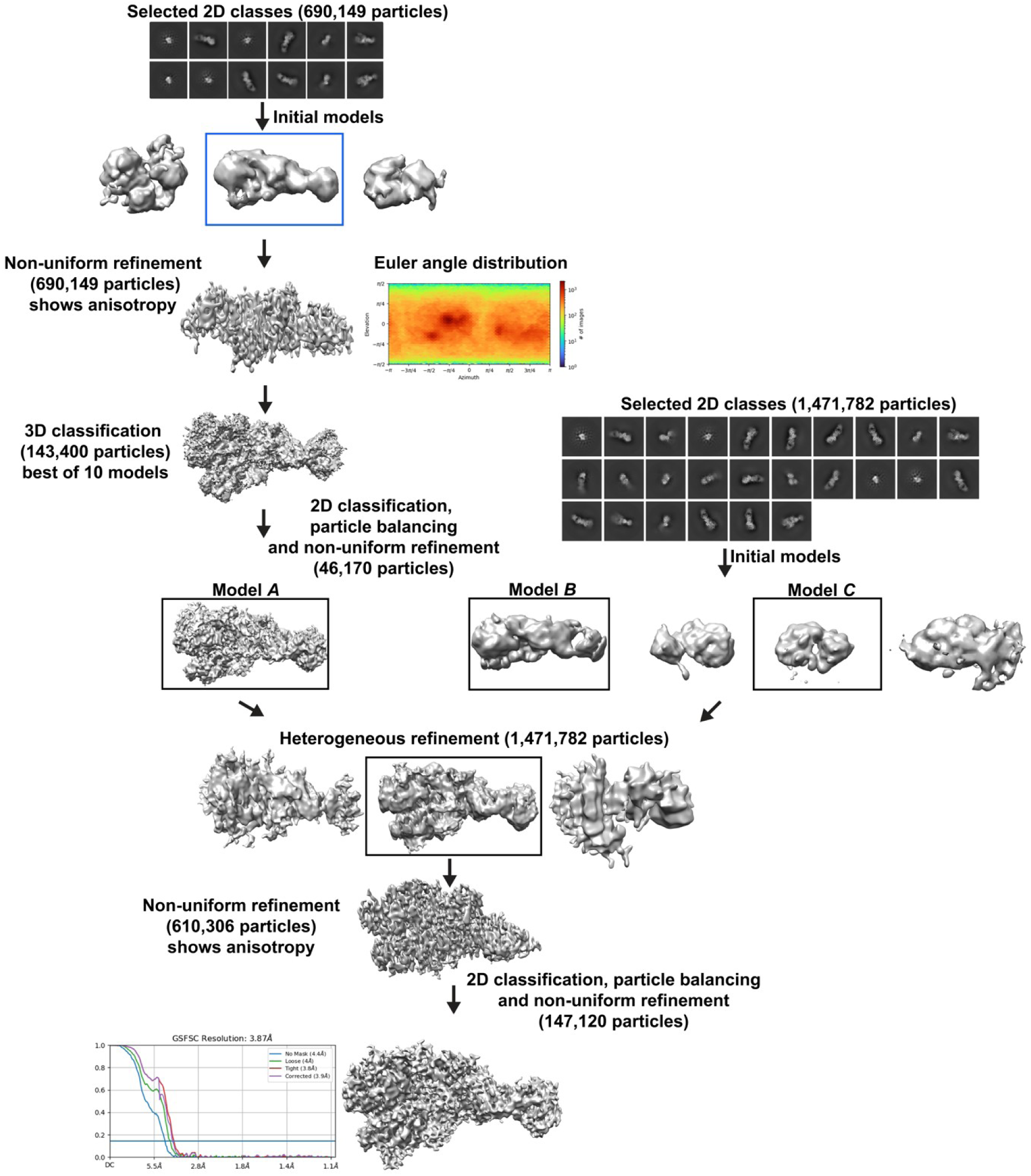
Cryo-EM workflow. Twelve 2D classes with density for Fab2 or PLCε PH-C and/or secondary structure features were selected and used to generate three *ab initio* models. The best model (boxed in blue) was used for a non-uniform refinement. However, the resulting map and Euler angle analysis revealed anisotropy consistent with a preferred orientation, likely due to Fab2. To address this, model *A* was subjected to 3D classification, 2D re-selection, and particle balancing. To increase the total number of particles (46,170 after re-selection and balancing), more 2D classes were selected (1,471,782 particles) and used to generate four *ab initio* models. Two of the four classes were consistent with Fab2– PLCε PH-C (model *B*) and Fab2 alone (model *C*). Heterogeneous refinement was performed to separate the 1,471,782 particles using these models. The class showing clear density for Fab2–PLCε PH-C was then used for a non-uniform refinement with decreased anisotropy. This was further corrected using 2D classification and particle balancing.

**Supporting Figure 3.**
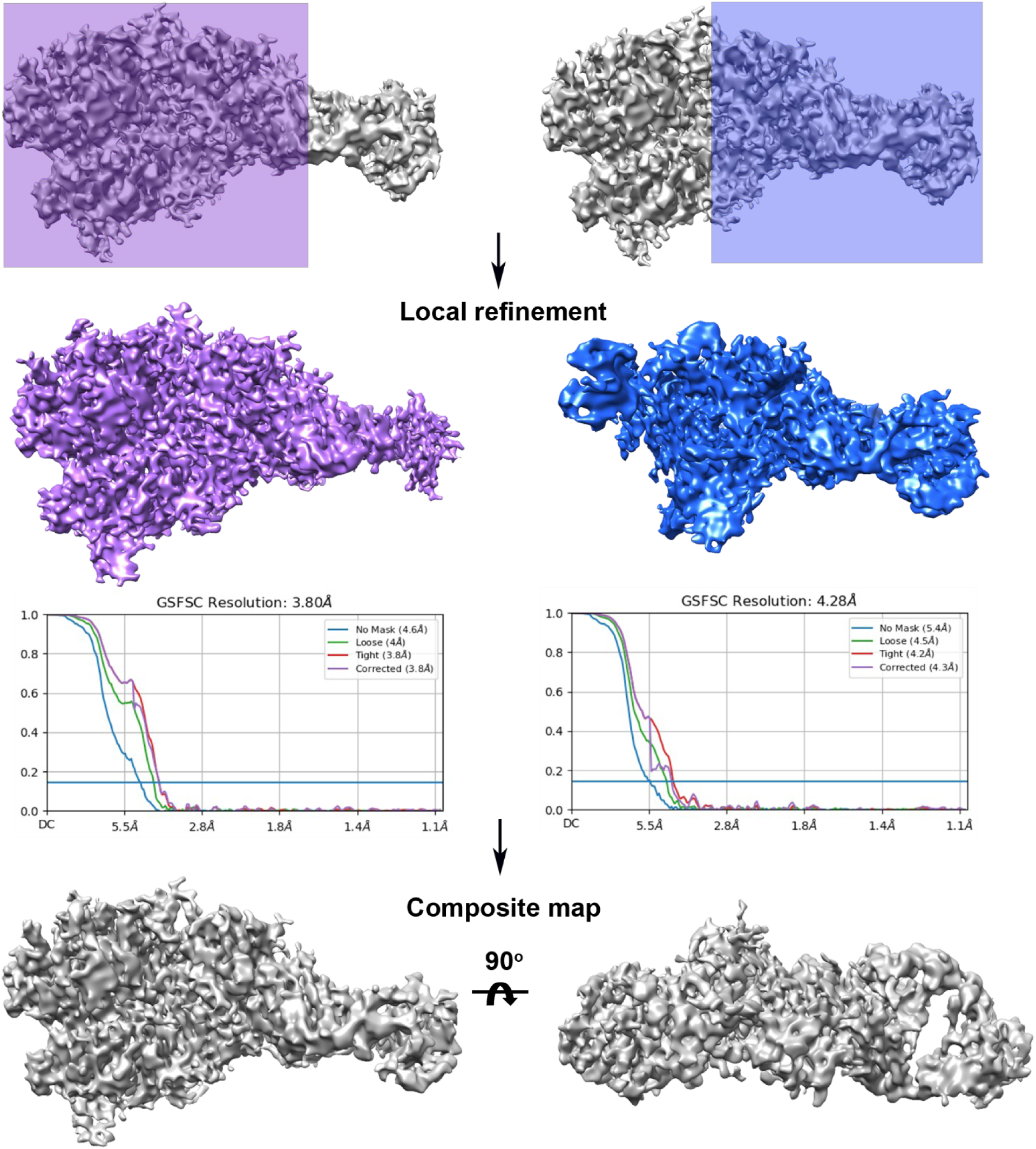
Generation of a composite map for Fab2–PLCε PH-C. The density for PLCε PH-C and Fab2 were improved by local refinement using different masks on cryoSPARC^23^. Shaded areas on the non-uniform map indicate the regions that were locally refined, including PLCε PH-C (purple) and the Fab2–PLCε PH-C interface (blue). The non-uniform, locally refined maps were then used to generate a final composite map at 3.9 Å resolution using PHENIX^48^.

**Supporting Figure 4.**
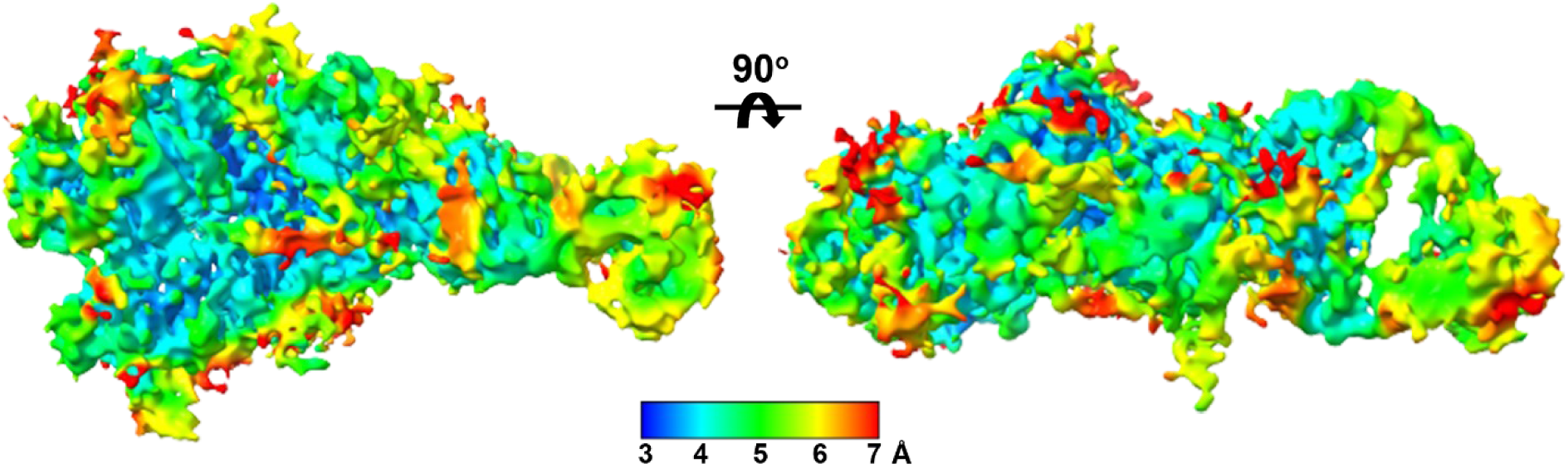
Resolution estimate of the Fab2–PLCε PH-C complex composite map. Cryo-EM final composite map of the Fab2–PLCε PH-C complex colored according to local resolution calculated by cryoSPARC^23^. The numbers on the scale bar correspond to the resolution in Å.

**Supporting Figure 5.**
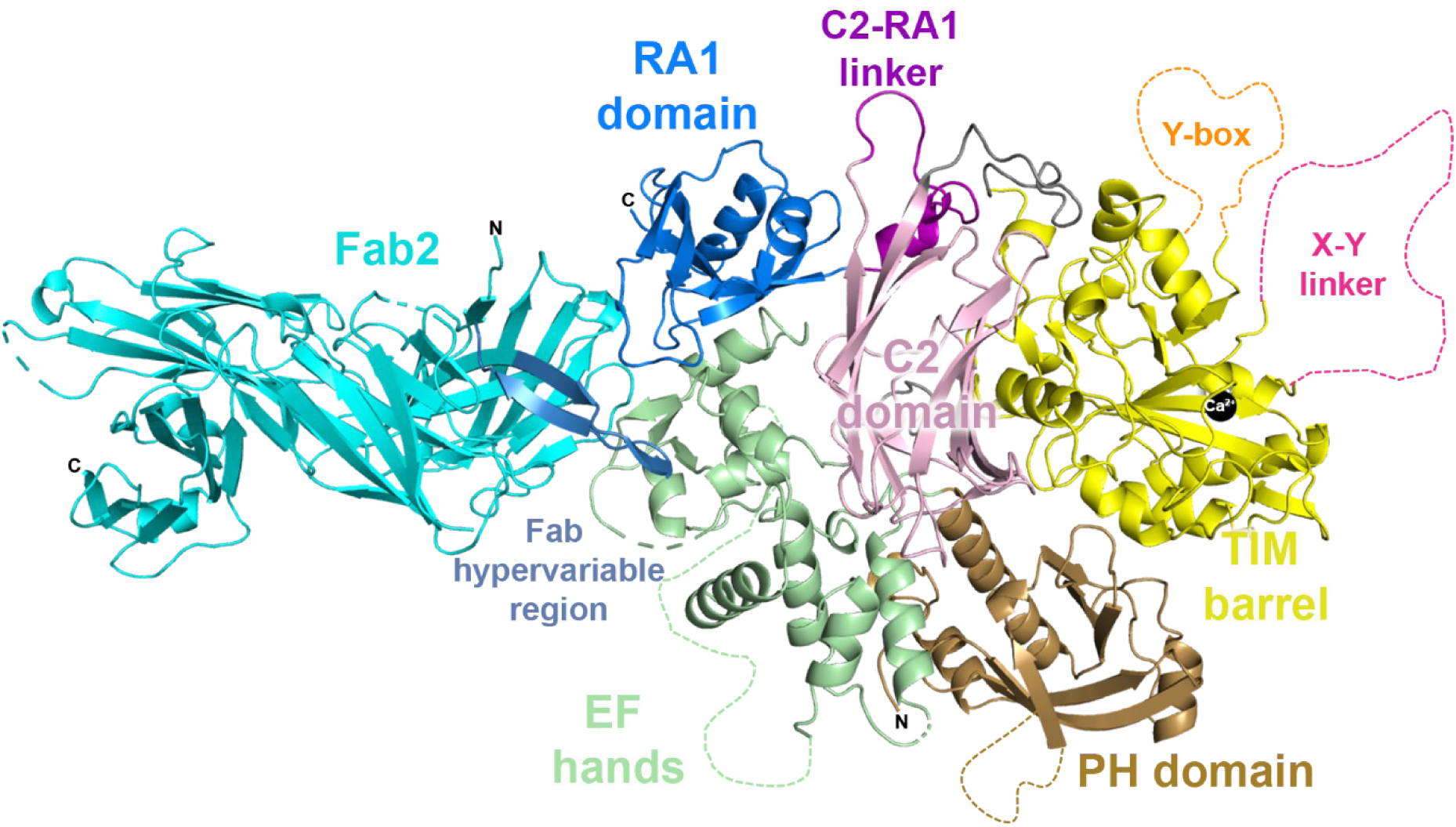
Fab2 binds the EF hands and RA1 domains. The hypervariable region of Fab2 interacts directly with the EF hands and the RA1 domain, burying ∼2,200 Å^2^ surface area.

**Supporting Figure 6.**
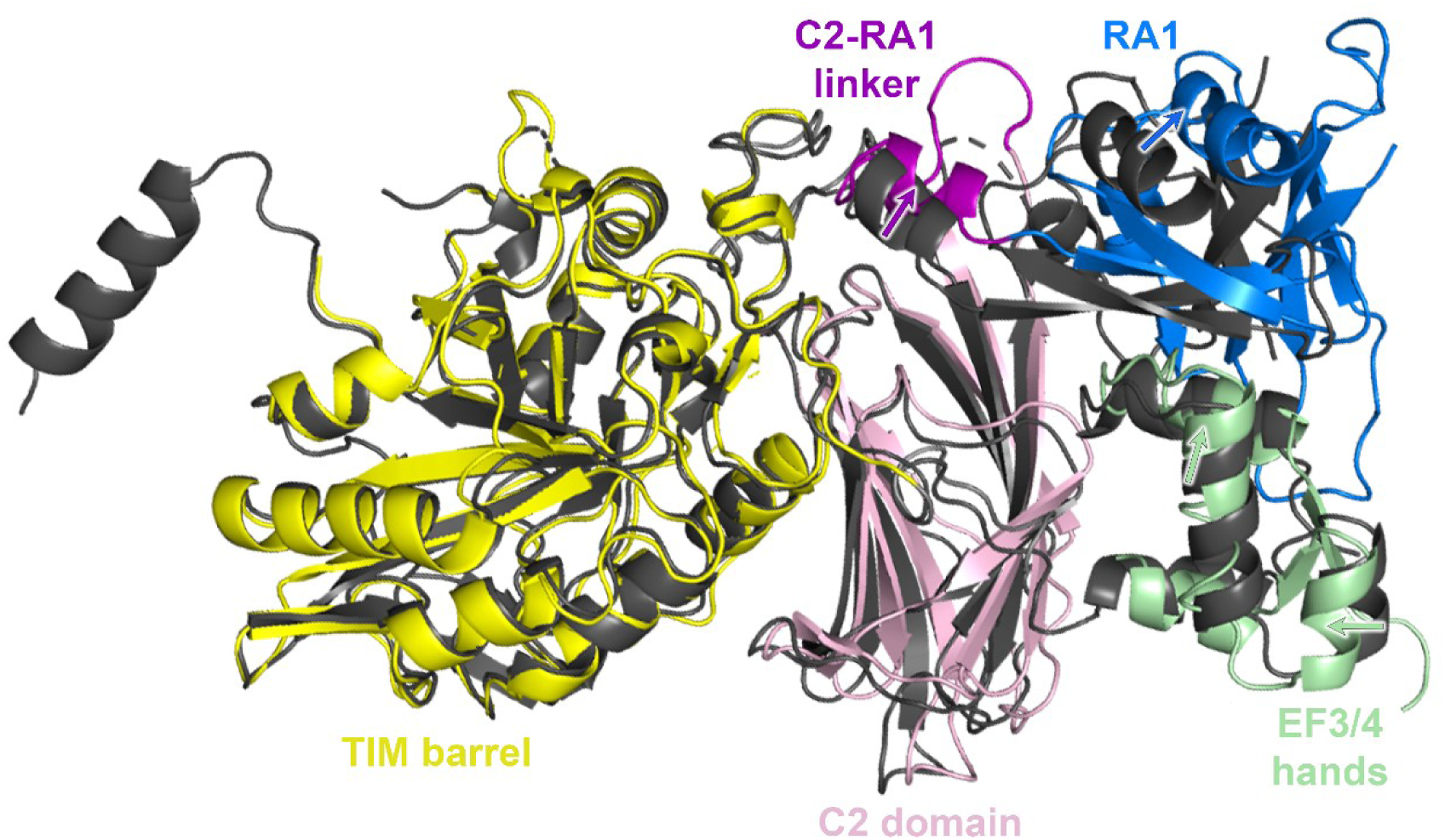
Conformational differences in the PLCε EF3-RA1 catalytic core in solution. The crystal structure of EF3-RA1 (dark grey, PDB ID: 6PMP)^13^ and the reconstruction of the Fab2**–**PLCε PH-C complex were compared using residue-residue (RR) distance maps^49^ in ChimeraX^50^, revealing the position of the TIM barrel between structures was nearly identical. The crystal structure and cryo-EM reconstruction were superimposed over the TIM barrel to identify any changes in other regions of the protein. Relative to their positions in the crystal structure, the RA1 domain is shifted ∼7 Å away from the TIM barrel in the cryo-EM reconstruction, as is the C2-RA1 linker. EF3/4 is also shifted ∼4 Å from its position in the crystal structure towards the C2 domain. Minor shifts in the positions of the loops in the C2 domain that interact with the TIM barrel and EF3/4 hands are also observed.

**Supporting Figure 7.**
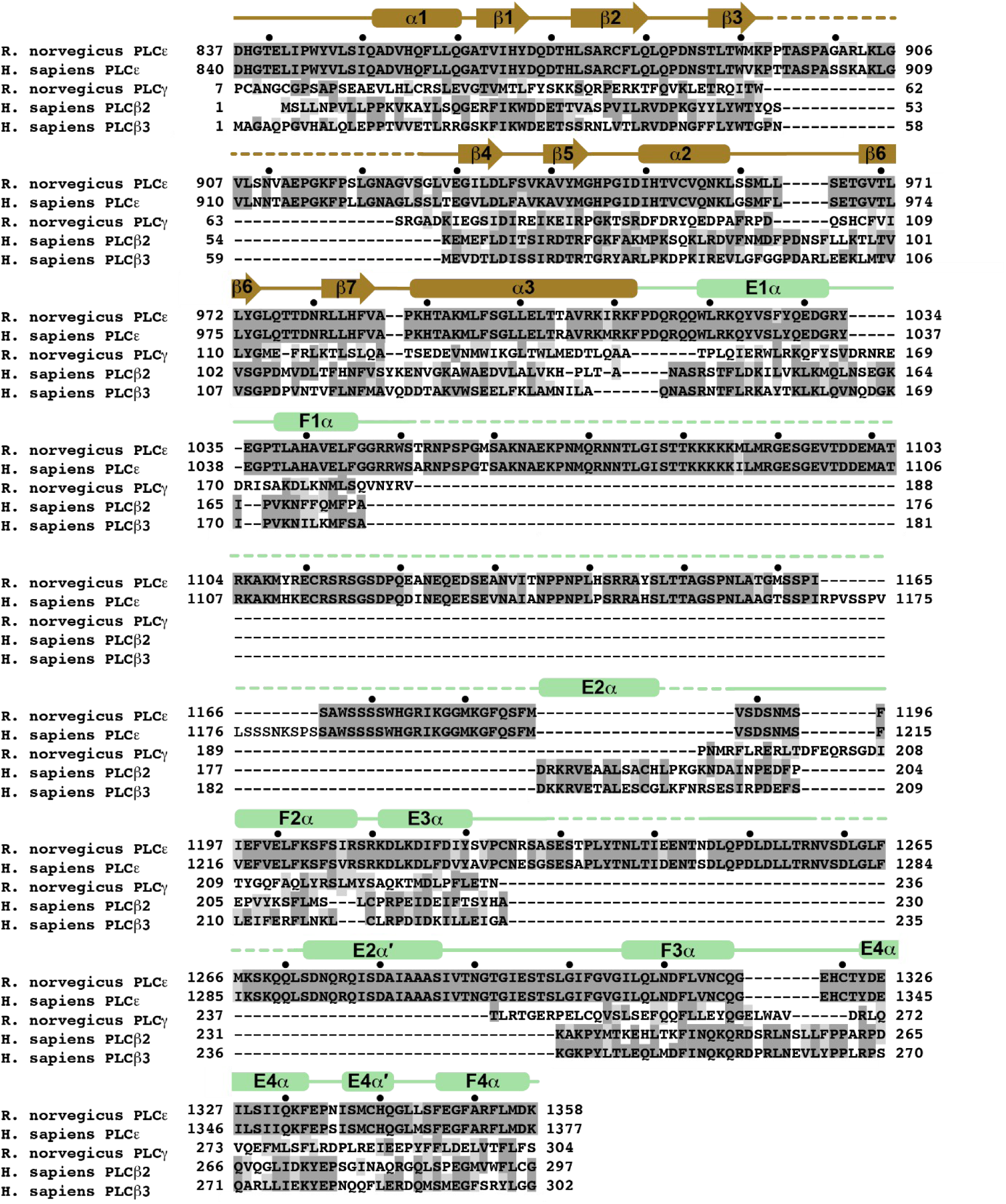
Alignment of the PLC PH and EF hand domains. The amino acid sequences of *R. norvegicus* PLCε (UNIPROT Q99P84), *H. sapiens*; PLCε (UNIPROT Q9P212), *R. norvegicus* PLCγ1 (UNIPROT P10686), *H. sapiens* PLCβ2 (UNIPROT Q00722), *H. sapiens* PLCβ3 (UNIPROT Q01970) PH domains (brown) and EF hands (green) are shown. Black circles above are spaced ten residues apart in *R. norvegicus* PLCε. Identical residues are highlighted in dark gray and similar residues in light gray. Observed secondary structure is shown above the alignment, with β strands shown as arrows, α helices as rounded rectangles, and dashed lines as disordered regions.

**Supporting Figure 8.**
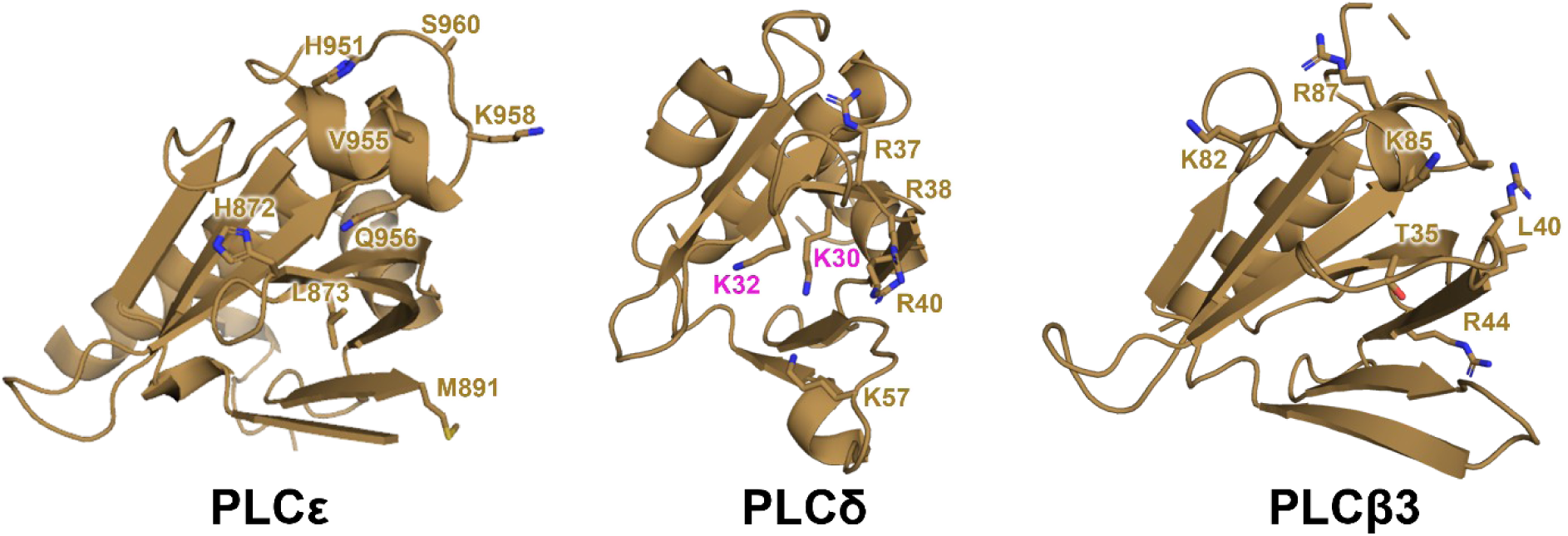
Comparison of PLC PH domains. The PH domains of PLCε, PLCδ (PDB ID: 1MAI)^51^, and PLCβ3 (PDB ID: 3OHM)^25^ are oriented to view their membrane binding surfaces. The structures are similar, with an overall r.m.s.d. of 1-1.14 Å for 106 Cα atoms. Residues required for PIP_2_ binding in PLCδ are labelled in magenta. These residues are not conserved in PLCε or PLCβ.

**Supporting Figure 9.**
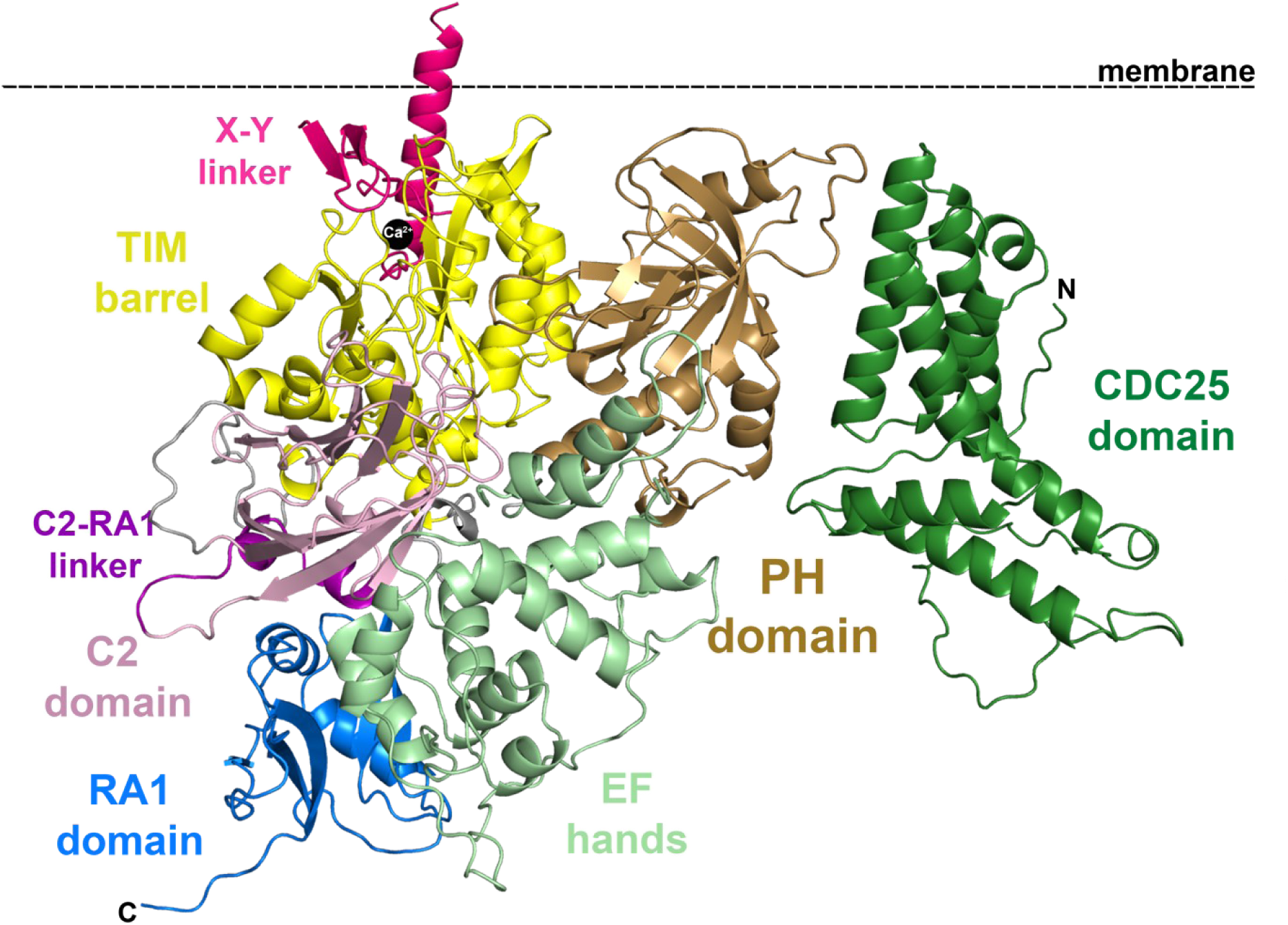
AlphaFold2^17^ model of PLCε CDC25-RA1. The AlphaFold2^17^ model predicts an interface between the CDC25 (green) and PH domain (wheat) that buries ∼1,400 Å^2^ surface area. The basic and hydrophobic surface formed by the two domains is in the same plane as the active site in the TIM barrel, denoted by the catalytic Ca^2+^ ion (black sphere). Dashed lines represent disordered loops, and the N-and C-termini are labelled N and C, respectively.

**Table S2.**
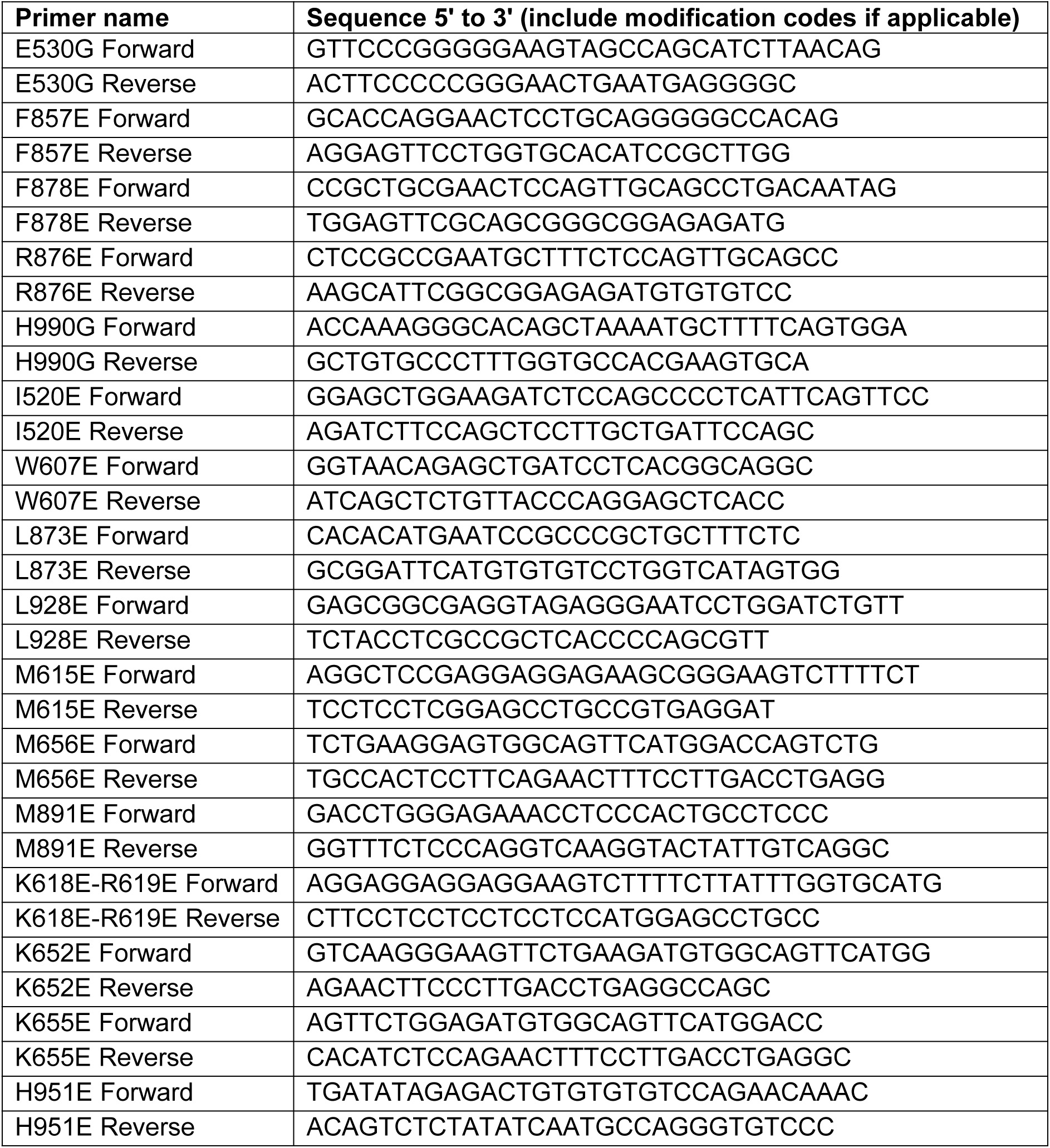
Primers used for site-directed mutagenesis.

## REFERENCES

1. Kadamur, G.; Ross, E. M., Mammalian Phospholipase C. Annual Review of Physiology 2013, 75 (1), 127–154.

2. Muralidharan, K.; Van Camp, M. M.; Lyon, A. M., Structure and regulation of phospholipase Cβ and ε at the membrane. Chem Phys Lipids 2021, 235, 105050.

3. Wing, M. R.; Bourdon Dm Fau -Harden, T. K.; Harden, T. K., PLC-epsilon: a shared effector protein in Ras-, Rho-, and G alpha beta gamma-mediated signaling. (1534-0384 (Print)).

4. Smrcka, A. V.; Brown, J. H.; Holz, G. G., Role of phospholipase Cε in physiological phosphoinositide signaling networks. Cellular signalling 2012, 24 (6), 1333–1343.

5. Madukwe, J. C.; Garland-Kuntz, E. E.; Lyon, A. M.; Smrcka, A. V., G protein βγ subunits directly interact with and activate phospholipase Cɛ. (1083–351X (Electronic)).

6. Zhang, L.; Malik, S.; Kelley, G. G.; Kapiloff, M. S.; Smrcka, A. V., Phospholipase C epsilon scaffolds to muscle-specific A kinase anchoring protein (mAKAPbeta) and integrates multiple hypertrophic stimuli in cardiac myocytes. The Journal of biological chemistry 2011, 286 (26), 23012–23021.

7. Nash, C. A.; Wei, W.; Irannejad, R.; Smrcka, A. V., Golgi localized β1-adrenergic receptors stimulate Golgi PI4P hydrolysis by PLCε to regulate cardiac hypertrophy. eLife 2019, 8, e48167.

8. Jin, T. G.; Satoh T Fau - Liao, Y.; Liao Y Fau - Song, C.; Song C Fau - Gao, X.; Gao X Fau - Kariya, K.; Kariya K Fau - Hu, C. D.; Hu Cd Fau - Kataoka, T.; Kataoka, T., Role of the CDC25 homology domain of phospholipase Cepsilon in amplification of Rap1-dependent signaling. (0021–9258 (Print)).

9. Sieng, M.; Selvia, A. F.; Garland-Kuntz, E. E.; Hopkins, J. B.; Fisher, I. J.; Marti, A. T.; Lyon, A. M., Functional and structural characterization of allosteric activation of phospholipase Cε by Rap1A. J Biol Chem 2020, 295 (49), 16562–16571.

10. Wang, H.; Oestreich Ea Fau - Maekawa, N.; Maekawa N Fau - Bullard, T. A.; Bullard Ta Fau - Vikstrom, K. L.; Vikstrom Kl Fau - Dirksen, R. T.; Dirksen Rt Fau - Kelley, G. G.; Kelley Gg Fau - Blaxall, B. C.; Blaxall Bc Fau - Smrcka, A. V.; Smrcka, A. V., Phospholipase C epsilon modulates beta-adrenergic receptor-dependent cardiac contraction and inhibits cardiac hypertrophy. (1524-4571 (Electronic)).

11. Xiang, S. Y.; Vanhoutte D Fau - Del Re, D. P.; Del Re Dp Fau - Purcell, N. H.; Purcell Nh Fau - Ling, H.; Ling H Fau - Banerjee, I.; Banerjee I Fau - Bossuyt, J.; Bossuyt J Fau - Lang, R. A.; Lang Ra Fau - Zheng, Y.; Zheng Y Fau - Matkovich, S. J.; Matkovich Sj Fau - Miyamoto, S.; Miyamoto S Fau - Molkentin, J. D.; Molkentin Jd Fau - Dorn, G. W., 2nd; Dorn Gw 2nd Fau - Brown, J. H.; Brown, J. H., RhoA protects the mouse heart against ischemia/reperfusion injury. (1558-8238 (Electronic)).

12. Xiang, S. Y.; Ouyang K Fau - Yung, B. S.; Yung Bs Fau - Miyamoto, S.; Miyamoto S Fau - Smrcka, A. V.; Smrcka Av Fau - Chen, J.; Chen J Fau - Heller Brown, J.; Heller Brown, J., PLCε, PKD1, and SSH1L transduce RhoA signaling to protect mitochondria from oxidative stress in the heart. (1937-9145 (Electronic)).

13. Rugema, N. Y.; Garland-Kuntz, E. E.; Sieng, M.; Muralidharan, K.; Van Camp, M. M.; O’Neill, H.; Mbongo, W.; Selvia, A. F.; Marti, A. T.; Everly, A.; McKenzie, E.; Lyon, A. M., Structure of phospholipase Cε reveals an integrated RA1 domain and previously unidentified regulatory elements. Commun Biol 2020, 3 (1), 445.

14. Song, C.; Hu, C.-D.; Masago, M.; Kariya, K.-i.; Yamawaki-Kataoka, Y.; Shibatohge, M.; Wu, D.; Satoh, T.; Kataoka, T., Regulation of a Novel Human Phospholipase C, PLCε, through Membrane Targeting by Ras *. Journal of Biological Chemistry 2001, 276 (4), 2752–2757.

15. Garland-Kuntz, E. E.; Vago, F. S.; Sieng, M.; Van Camp, M.; Chakravarthy, S.; Blaine, A.; Corpstein, C.; Jiang, W.; Lyon, A. M., Direct observation of conformational dynamics of the PH domain in phospholipases Cɛ and β may contribute to subfamily-specific roles in regulation. The Journal of biological chemistry 2018, 293 (45), 17477–17490.

16. Bunney, T. D.; Harris R Fau - Gandarillas, N. L.; Gandarillas Nl Fau - Josephs, M. B.; Josephs Mb Fau - Roe, S. M.; Roe Sm Fau - Sorli, S. C.; Sorli Sc Fau - Paterson, H. F.; Paterson Hf Fau - Rodrigues-Lima, F.; Rodrigues-Lima F Fau - Esposito, D.; Esposito D Fau - Ponting, C. P.; Ponting Cp Fau - Gierschik, P.; Gierschik P Fau - Pearl, L. H.; Pearl Lh Fau - Driscoll, P. C.; Driscoll Pc Fau - Katan, M.; Katan, M., Structural and mechanistic insights into ras association domains of phospholipase C epsilon. (1097-2765 (Print)).

17. Jumper, J.; Evans, R.; Pritzel, A.; Green, T.; Figurnov, M.; Ronneberger, O.; Tunyasuvunakool, K.; Bates, R.; Žídek, A.; Potapenko, A.; Bridgland, A.; Meyer, C.; Kohl, S. A. A.; Ballard, A. J.; Cowie, A.; Romera-Paredes, B.; Nikolov, S.; Jain, R.; Adler, J.; Back, T.; Petersen, S.; Reiman, D.; Clancy, E.; Zielinski, M.; Steinegger, M.; Pacholska, M.; Berghammer, T.; Bodenstein, S.; Silver, D.; Vinyals, O.; Senior, A. W.; Kavukcuoglu, K.; Kohli, P.; Hassabis, D., Highly accurate protein structure prediction with AlphaFold. Nature 2021, 596 (7873), 583–589.

18. Davydova, E. K., Protein Engineering: Advances in Phage Display for Basic Science and Medical Research. (1608-3040 (Electronic)).

19. Ereño-Orbea, J.; Sicard, T.; Cui, H.; Carson, J.; Hermans, P.; Julien, J. P., Structural Basis of Enhanced Crystallizability Induced by a Molecular Chaperone for Antibody Antigen-Binding Fragments. (1089-8638 (Electronic)).

20. Dutka, P.; Mukherjee, S.; Gao, X.; Kang, Y.; de Waal, P. W.; Wang, L.; Zhuang, Y.; Melcher, K.; Zhang, C.; Xu, H. E.; Kossiakoff, A. A., Development of “Plug and Play” Fiducial Marks for Structural Studies of GPCR Signaling Complexes by Single-Particle Cryo-EM. Structure 2019, 27 (12), 1862–1874.e7.

21. Bloch, J. A.-O.; Mukherjee, S. A.-O.; Kowal, J. A.-O.; Filippova, E. A.-O.; Niederer, M.; Pardon, E.; Steyaert, J.; Kossiakoff, A. A.; Locher, K. A.-O., Development of a universal nanobody-binding Fab module for fiducial-assisted cryo-EM studies of membrane proteins. LID - 10.1073/pnas.2115435118 [doi] LID - e2115435118. (1091-6490 (Electronic)).

22. Miller, K. R.; Koide, A.; Leung, B.; Fitzsimmons, J.; Yoder, B.; Yuan, H.; Jay, M.; Sidhu, S. S.; Koide, S.; Collins, E. J., T Cell Receptor-Like Recognition of Tumor In Vivo by Synthetic Antibody Fragment. PLOS ONE 2012, 7 (8), e43746.

23. Punjani, A.; Rubinstein, J. L.; Fleet, D. J.; Brubaker, M. A., cryoSPARC: algorithms for rapid unsupervised cryo-EM structure determination. Nat Methods 2017, 14 (3), 290–296.

24. Mukherjee, S.; Griffin, D. H.; Horn, J. R.; Rizk, S. S.; Nocula-Lugowska, M.; Malmqvist, M.; Kim, S. S.; Kossiakoff, A. A., Engineered synthetic antibodies as probes to quantify the energetic contributions of ligand binding to conformational changes in proteins. Journal of Biological Chemistry 2018, 293 (8), 2815–2828.

25. Waldo, G. L.; Ricks, T. K.; Hicks, S. N.; Cheever, M. L.; Kawano, T.; Tsuboi, K.; Wang, X.; Montell, C.; Kozasa, T.; Sondek, J.; Harden, T. K., Kinetic Scaffolding Mediated by a Phospholipase C– β and Gq Signaling Complex. Science 2010, 330 (6006), 974–980.

26. Garcia, P.; Gupta R Fau - Shah, S.; Shah S Fau - Morris, A. J.; Morris Aj Fau - Rudge, S. A.; Rudge Sa Fau - Scarlata, S.; Scarlata S Fau - Petrova, V.; Petrova V Fau - McLaughlin, S.; McLaughlin S Fau - Rebecchi, M. J.; Rebecchi, M. J., The pleckstrin homology domain of phospholipase C-delta 1 binds with high affinity to phosphatidylinositol 4,5-bisphosphate in bilayer membranes. (0006-2960 (Print)).

27. Lemmon, M. A.; Ferguson Km Fau - O’Brien, R.; O’Brien R Fau - Sigler, P. B.; Sigler Pb Fau - Schlessinger, J.; Schlessinger, J., Specific and high-affinity binding of inositol phosphates to an isolated pleckstrin homology domain. (0027-8424 (Print)).

28. Yagisawa, H.; Sakuma K Fau - Paterson, H. F.; Paterson Hf Fau - Cheung, R.; Cheung R Fau - Allen, V.; Allen V Fau - Hirata, H.; Hirata H Fau - Watanabe, Y.; Watanabe Y Fau - Hirata, M.; Hirata M Fau - Williams, R. L.; Williams Rl Fau - Katan, M.; Katan, M., Replacements of single basic amino acids in the pleckstrin homology domain of phospholipase C-delta1 alter the ligand binding, phospholipase activity, and interaction with the plasma membrane. (0021-9258 (Print)).

29. Ferguson, K. M.; Lemmon Ma Fau - Schlessinger, J.; Schlessinger J Fau - Sigler, P. B.; Sigler, P. B., Structure of the high affinity complex of inositol trisphosphate with a phospholipase C pleckstrin homology domain. (0092-8674 (Print)).

30. Tall, E.; Dormán G Fau - Garcia, P.; Garcia P Fau - Runnels, L.; Runnels L Fau - Shah, S.; Shah S Fau - Chen, J.; Chen J Fau - Profit, A.; Profit A Fau - Gu, Q. M.; Gu Qm Fau - Chaudhary, A.; Chaudhary A Fau - Prestwich, G. D.; Prestwich Gd Fau - Rebecchi, M. J.; Rebecchi, M. J., Phosphoinositide binding specificity among phospholipase C isozymes as determined by photo-cross-linking to novel substrate and product analogs. (0006-2960 (Print)).

31. Wang, T.; Pentyala S Fau - Rebecchi, M. J.; Rebecchi Mj Fau - Scarlata, S.; Scarlata, S., Differential association of the pleckstrin homology domains of phospholipases C-beta 1, C-beta 2, and C-delta 1 with lipid bilayers and the beta gamma subunits of heterotrimeric G proteins. (0006-2960 (Print)).

32. Philip, F.; Guo Y Fau - Scarlata, S.; Scarlata, S., Multiple roles of pleckstrin homology domains in phospholipase Cbeta function. (0014-5793 (Print)).

33. Lyon, A. M.; Tesmer, J. J., Structural insights into phospholipase C-β function. Mol Pharmacol 2013, 84 (4), 488–500.

34. Rebecchi, M. J.; Pentyala, S. N., Structure, Function, and Control of Phosphoinositide-Specific Phospholipase C. Physiological Reviews 2000, 80 (4), 1291–1335.

35. Essen, L. O.; Perisic O Fau - Cheung, R.; Cheung R Fau - Katan, M.; Katan M Fau - Williams, R. L.; Williams, R. L., Crystal structure of a mammalian phosphoinositide-specific phospholipase C delta. (0028-0836 (Print)).

36. Seifert, J. P.; Zhou Y Fau - Hicks, S. N.; Hicks Sn Fau - Sondek, J.; Sondek J Fau - Harden, T. K.; Harden, T. K., Dual activation of phospholipase C-epsilon by Rho and Ras GTPases. (0021-9258 (Print)).

37. Boriack-Sjodin, P. A.; Margarit, S. M.; Bar-Sagi, D.; Kuriyan, J., The structural basis of the activation of Ras by Sos. Nature 1998, 394 (6691), 337–343.

38. Zheng, J.; Chen, R. H.; Corblan-Garcia, S.; Cahill, S. M.; Bar-Sagi, D.; Cowburn, D., The Solution Structure of the Pleckstrin Homology Domain of Human SOS1: A POSSIBLE STRUCTURAL ROLE FOR THE SEQUENTIAL ASSOCIATION OF DIFFUSE B CELL LYMPHOMA AND PLECKSTRIN HOMOLOGY DOMAINS *. Journal of Biological Chemistry 1997, 272 (48), 30340–30344.

39. Kelley, G. G.; Reks Se Fau - Smrcka, A. V.; Smrcka, A. V., Hormonal regulation of phospholipase Cepsilon through distinct and overlapping pathways involving G12 and Ras family G-proteins. (1470-8728 (Electronic)).

40. Paduch, M.; Kossiakoff, A. A., Generating Conformation and Complex-Specific Synthetic Antibodies. In Synthetic Antibodies: Methods and Protocols, Tiller, T., Ed. Springer New York: New York, NY, 2017; pp 93–119.

41. Hornsby, M.; Paduch, M.; Miersch, S.; Sääf, A.; Matsuguchi, T.; Lee, B.; Wypisniak, K.; Doak, A.; King, D.; Usatyuk, S.; Perry, K.; Lu, V.; Thomas, W.; Luke, J.; Goodman, J.; Hoey, R. J.; Lai, D.; Griffin, C.; Li, Z.; Vizeacoumar, F. J.; Dong, D.; Campbell, E.; Anderson, S.; Zhong, N.; Gräslund, S.; Koide, S.; Moffat, J.; Sidhu, S.; Kossiakoff, A.; Wells, J., A High Through-put Platform for Recombinant Antibodies to Folded Proteins. (1535-9484 (Electronic)).

42. Yu, Y.; Zheng, Q.; Erramilli, S. A.-O.; Pan, M. A.-O.; Park, S.; Xie, Y. A.-O.; Li, J.; Fei, J. A.-O.; Kossiakoff, A. A.; Liu, L. A.-O.; Zhao, M. A.-O., K29-linked ubiquitin signaling regulates proteotoxic stress response and cell cycle. (1552-4469 (Electronic)).

43. Liebschner, D.; Afonine, P. V.; Baker, M. L.; Bunkóczi, G.; Chen, V. B.; Croll, T. I.; Hintze, B.; Hung, L.-W.; Jain, S.; McCoy, A. J.; Moriarty, N. W.; Oeffner, R. D.; Poon, B. K.; Prisant, M. G.; Read, R. J.; Richardson, J. S.; Richardson, D. C.; Sammito, M. D.; Sobolev, O. V.; Stockwell, D. H.; Terwilliger, T. C.; Urzhumtsev, A. G.; Videau, L. L.; Williams, C. J.; Adams, P. D., Macromolecular structure determination using X-rays, neutrons and electrons: recent developments in Phenix. Acta Crystallographica Section D Structural Biology 2019, 75 (10), 861–877.

44. Yang, J.; Zhang, Y., Protein Structure and Function Prediction Using I-TASSER. (1934-340X (Electronic)).

45. Emsley, P.; Cowtan, K., Coot: model-building tools for molecular graphics. (0907-4449 (Print)).

46. CisBio, IP-One Gq kit.

47. Esquina, C. M.; Garland-Kuntz, E. E.; Goldfarb, D.; McDonald, E. K.; Hudson, B. N.; Lyon, A. M., Intramolecular electrostatic interactions contribute to phospholipase Cβ3 autoinhibition. Cell Signal 2019, 62, 109349.

48. Liebschner, D. A.-O.; Afonine, P. A.-O. X.; Baker, M. L.; Bunkóczi, G.; Chen, V. B.; Croll, T. I.; Hintze, B. A.-O.; Hung, L. A.-O.; Jain, S.; McCoy, A. J.; Moriarty, N. A.-O.; Oeffner, R. A.-O.; Poon, B. A.-O.; Prisant, M. G.; Read, R. A.-O.; Richardson, J. S.; Richardson, D. C.; Sammito, M. D.; Sobolev, O. A.-O.; Stockwell, D. H.; Terwilliger, T. A.-O.; Urzhumtsev, A. G.; Videau, L. L.; Williams, C. J.; Adams, P. A.-O., Macromolecular structure determination using X-rays, neutrons and electrons: recent developments in Phenix. (2059-7983 (Electronic)).

49. Schneider, T. R., Objective comparison of protein structures: error-scaled difference distance matrices. (0907-4449 (Print)).

50. Meng, E. C.; Goddard, T. D.; Pettersen, E. F.; Couch, G. S.; Pearson, Z. J.; Morris, J. H.; Ferrin, T. E., UCSF ChimeraX: Tools for structure building and analysis. Protein Science 2023, 32 (11), e4792.

51. Ferguson, K. M.; Lemmon, M. A.; Schlessinger, J.; Sigler, P. B., Structure of the high affinity complex of inositol trisphosphate with a phospholipase C pleckstrin homology domain. Cell 1995, 83 (6), 1037–1046.

